# Noradrenergic neuromodulation produces a NMDAR-dependent network state of respiratory rhythmogenesis in the preBötzinger Complex

**DOI:** 10.64898/2026.02.11.705209

**Authors:** Andrew Kieran Tryba, Jean-Charles Viemari, Yangyang Wang, Alfredo J. Garcia

**Affiliations:** Department of Pediatrics, Section of Neurology, The University of Chicago, Chicago, IL, 60637, United States of America; Institute for Integrative Physiology, The University of Chicago, Chicago, IL, 60637, United States of America; Marseille Medical Genetics, UMR1251, Aix Marseille University, Inserm, 27 Bd Jean Moulin 13005 Marseille, France; INT, UMR 7289, AMU-CNRS, 27 Bd Jean Moulin 13005 Marseille, France; Department of Mathematics, Neuroscience Program, Volen National Center for Complex Systems, Brandeis University, Waltham, MA, 02453-2728, United States of America; Department of Medicine, Section of Emergency Medicine, The University of Chicago, Chicago, IL, 60637, United States of America; The University of Chicago Neuroscience Institute, The University of Chicago, Chicago, IL, 60637, United States of America

## Abstract

Norepinephrine (NE) is an important mediator of sympathetic activity that influences breathing. At the level of the inspiratory neural network, the preBötzinger complex (preBötC), NE modulation orchestrates changes in neuronal network dynamics that influence the stability of inspiratory rhythmogenesis. While this phenomenon has largely been attributed to NE-modulation of intrinsic excitability of inspiratory preBötC neurons, NE is also capable of modulating synaptic drive. Here, we resolve how NE affects synaptic properties and changes the activity dynamics of interconnected preBötC neurons in rhythmic brainstem slice preparations. Increased network burst amplitude and frequency coincided with enhanced inspiratory drive currents at the single neuron level. This increased drive was blocked by the NMDA receptor (NMDAR) antagonist, APV. Our *in silico* modeling indicated that synaptic calcium entry via NDMAR is key to maintaining network synchrony during this elevated state of excitability. This was consistent with our multi-electrode array studies revealing that NE-dependent NMDAR activity enhances and preserves synchrony during inspiratory network bursts. This synaptic mechanism may be a critical determinant for shaping inspiratory drive associated with changed neuromodulatory environments.

**Significance Statement:** This study demonstrates a previously undescribed synaptic mechanism by which noradrenergic modulation recruits NMDAR activity and shifts inspiratory network dynamics leading to a stable network state where synchronization is preserved among inspiratory neurons. This may be a critical mechanism preventing ventilatory instability when sympathetic activity is enhanced.

## Introduction

Coordinated activity is a fundamental property of neural networks that supports the neurophysiological processes of cognition, memory and vital autonomic functions such as breathing. With respect to respiratory control, the preBötzinger Complex (preBötC) serves as the kernel for inspiratory rhythmogenesis, where the network rhythm emerges from coherent synchronized activity across a heterogeneous neuronal population. Efforts toward understanding the process of inspiratory rhythmogenesis established that synchronization is critical to both the generation and maintenance of the inspiratory rhythm and a loss of synchrony can lead to subnetwork activity that fails to generate downstream motor output at the level of the hypoglossal motor nucleus (XIIn) (Butera et al. 1999; Kam et al. 2013; Ashhad and Feldman 2020). Reducing excitability in *in vitro* rhythmic respiratory slice system reveals that such subnetworks can emerge in the preBötC and has led to the proposal of the so-called “burstlet” hypothesis, in which low amplitude rhythmogenic “burstlets” must be amplified into a suprathreshold “burst” to generate XIIn motor output (Ashhad and Feldman, 2020). While this process is proposed to be governed by the network level refractory period (Ashhad et al. 2023) and shaped by dynamic interactions preserving excitatory-inhibitory (E/I) synaptic balance within the preBötC (Harris et al. 2017; Ashhad and Feldman, 2020; Ashhad et al. 2023), neuromodulation also plays a crucial role in shaping network excitability in the preBötC.

Neuromodulation of the inspiratory network plays a central role in allowing the network to respond to homeostatic challenges, such as hypoxia (Tryba et al. 2006; Doi and Ramirez 2008; Viemari et al. 2011). Among these, norepinephrine (NE), is a potent neuromodulator of preBötC activity stimulating several aspects of rhythmogenesis (Viemari and Ramirez, 2006; Viemari et al., 2011; Viemari et al. 2013). Prior investigations have emphasized the context-dependent nature of NE modulation (Viemari and Ramirez, 2006, Viemari et al. 2011; Zanella et al. 2014; Venkatakrishnan et al. 2026 (*in press*)) where the ability to stabilize rhythmogenesis has been principally attributed to changes in intrinsic membrane currents, such as the non-specific cation current (ICAN) and persistent sodium current (INaP) (Viemari and Ramirez, 2006). However, this intrinsic-centric framework may be incomplete in terms of understanding the influence of neuromodulators on intrinsic and synaptic properties that alter network dynamics.

Recent work demonstrating that opioid neuromodulation of the preBötC simultaneously modulates intrinsic membrane conductances and synaptic communication (Baertsch et al. 2021) raise the possibility that this is a broader principle by which neuromodulators likely act to regulate network states rather than neuromodulation just simply scaling neuronal excitability. Indeed, while NE normally stabilizes preBötC network rhythms under baseline experimental conditions, following intermittent hypoxia exposure, subsequent NE modulation has a paradoxical effect and destabilizes preBötC inspiratory rhythmogenesis by generating subnetwork activity that fails to activate downstream motorneuron activity (Zanella et al. 2014). These maladaptive responses were hypothesized to involve disruption of the E/I balance within the network (Zanella et al. 2014), potentially due to the heterogeneous nature of synaptic connectivity among preBötC neurons (Kam et al. 2013). However, the synaptic mechanisms by which NE modulation shapes network dynamics in the preBötC network remains unresolved.

Using a combination of in vitro experiments and in silico modeling, we define how NE modulation of synaptic input impacts the preBötC network state. We demonstrate that NE alters network dynamics via a previously undescribed role of Ca^2+^ permeable NMDA receptors (NMDAR). The NE-modulated increase in NMDAR activity preserves neuronal synchronization during inspiratory bursts while simultaneously enhancing the intensity and frequency of preBötC activity in the NE-modulated state, revealing a novel pathway through which NE stabilizes and amplifies the inspiratory network rhythm.

## Methods

### Approval Statement

Experimental procedures were carried out in accordance with the United States Public Health Service and Institute for Laboratory Animal Research Guide for the Care and Use of Laboratory Animals. All animals were handled according to protocols approved by The University of Chicago Institutional Animal Care and Use Committee.

### Respiratory medullary brainstem preparation

All experiments used the transverse, rhythmic 650μm thick medullary brain-slice obtained from postnatal 8-13 day old (P8-P13) CD-1 outbred mice (Charles River Laboratories, Wilmington, MA)(Smith et al., 1991; Thoby-Brisson and Ramirez, 2001). CD-1 mice were anesthetized in isoflurane, then rapidly decapitated at the C3/C4 spinal level and the brain-stem was dissected in ice cold artificial cerebral spinal fluid (ACSF) that was equilibrated with carbogen (95% O2 and 5% CO2, pH=7.4). Rhythmic slice preparations containing the preBötC, were prepared by slicing the medulla using a vibratome (Leica, VT1000S, Nussloch, Germany) as described in detail elsewhere (Garcia et al., 2017). Briefly, the brainstem was glued onto an agar block with the rostral end up, mounted into a microtome and serially sliced from rostral to caudal until the rostral boundary of the preBötC was visible. This area is recognized by landmarks such as the inferior olive (IO), the nucleus ambiguous (NA) and the hypoglossal nucleus (XII). Brain slices (∼650μm thick) containing the preBötC were submerged in a recording chamber (6 mL) under circulating ACSF (flow rate 17 ml/min, total re-circulating volume = 200mL). As bath temperature can alter activity in the preBötC (Tryba and Ramirez, 2003; Tryba and Ramirez 2004), it was monitored and maintained at 30°C ± 0.7°C using a Warner Instrument Corp. (Hamden, CT, U.S.A.) TC-344B temperature regulator with an in-line solution heater (SH-27B).

### ACSF contained in mM

118 NaCl, 3 KCl, 1.5 CaCl2, 1 MgCl2*6H2O, 25 NaHCO3, 1 NaH2PO4 and 30 D-glucose, equilibrated with carbogen (95% O2 and 5% CO2, pH = 7.4). All ACSF chemicals were obtained from Sigma (St. Louis, MO). To obtain and maintain rhythmic population activity extracellular KCl was elevated from 3mM to 8mM over a span of 20 minutes and recordings commenced 10mins after 8mM [K^+^]o was introduced (Tryba et al. 2003).

### Electrophysiological recordings of preBötC activity

Single-electrode extracellular recordings were obtained with glass suction electrodes positioned on the surface of the respiratory brainstem slice on top of the preBötC (Tryba et al., 2003a). The extracellular signals were amplified 10,000-fold and filtered between 0.25 and 1.5 kHz using a pre-amplifier (built by JFIE electronics at The University of Chicago, Chicago, IL, U.S.A.) and a Model P-55 A.C. amplifier (Grass Instruments Technologies, Astro-Med, Inc., West Warwick RI, U.S.A.).

The preBötC network bursting is dominated by inspiratory neurons such that integrated preBötC (∫preBötC) network bursts are in-phase with integrated XII (∫XII) activity (Tryba et al., 2006). Thus, preBötC network bursts serve as a marker of fictive inspiration (Tryba et al., 2006; Tryba et al., 2008). PreBötC network activity (and in some experiments XII network data) was rectified and integrated using a custom built electronic integrator (JFI electronics, The University of Chicago, Chicago), with an integration time constant of 140ms. Both the filtered and integrated activity data were digitized with a Digidata acquisition system (Molecular Devices, CA) and stored on an IBM compatible PC using pClamp v10.1 (Molecular Devices, CA) software for off-line analysis.

To investigate the effects of NE on inspiratory drive currents and potentials, we used intracellular whole-cell voltage and current-clamp recordings obtained with a MultiClamp 700B amplifier (Molecular Devices, CA), applying the blind-patch technique to preBötC neurons in rhythmic respiratory brainstem slice preparations containing the preBötC network (Thoby-Brisson and Ramirez, 2001). Patch electrodes (3-5 MΩ) were manufactured from filamented borosilicate glass tubes (Clark G150F-4; Warner Instruments Corp., Hamden, CT, USA) and filled with an intracellular solution containing (in mM): 140 K-gluconic acid, 1 CaCl_2_*6H_2_O, 10 EGTA, 2 MgCl_2_*6H2O, 4 Na_2_ATP, 10 HEPES. Only inspiratory neurons active in-phase with the ∫preBötC network burst were recorded in current-and voltage-clamp studies. The discharge pattern of each inspiratory neuron was first identified in the cell-attached mode and remained similar in whole-cell configuration (Tryba et al., 2008). Current-clamp and voltage-clamp experiments were then performed in the whole cell patch-clamp mode. The Vm values were corrected for the liquid junction potential as calculated using pClamp v10 software (Molecular Devices, CA). For voltage clamp studies, giga-ohm (GΩ) seals were established and whole-cell currents were measured with an access resistance (Ra) of ≤25MΩ, corrected for fast and slow capacitive currents and series resistance compensated. Inherent in all similar studies, whole-cell voltage clamp recordings from neurons embedded in a functional network, including neurons in this study, are inherently subject to space-clamp issues that prevent measuring the magnitude of current changes with high precision, yet they do reflect relative changes in currents between treatments.

After identifying the lateral border of preBötC network activity using a single extracellular recording electrode placed on the surface of the slice, a thirty-two (32)-channel multi-electrode array (MEA)(catalog number: A1×32-Poly3-10mm-25s-17, NeuroNexus, Ann Arbor, MI) was inserted into the lateral border of the PreBötC using a remote-controlled micromanipulator (micromanipulator Model DC3001, World Precision Instruments (WPI), Sarasota, FL). The 32 channels of MEA data recordings were made using 15𝛍m diameter electrodes arranged in three columns, with 12 electrodes in the center column, 10 electrodes in both lateral columns and electrodes spaced 25𝛍m apart. The MEA was inserted within the preBötC, so that the electrodes were facing the lateral (rather than medial) region of the preBötC within the slice to minimize potential damage to the network and to neurons that decussate from the contralateral preBötC. The MEA was connected to a A32-Z32 adaptor (Tucker-Davis Technologies (TDT), Alachua, FL). The signal was amplified via a NeuroDigitizer PZ5-32 pre-amplifier (Tucker-Davis Technologies (TDT), Alachua, FL). Data was then recorded using a TDT WS8 computer workstation and TDT Synapse software (version 98-51640).

### Drugs

In some experiments, neurons were isolated from ionotropic chemical synaptic input using a mixture of glutamatergic, GABAergic and glycinergic antagonists. These drugs were bath-applied at the final concentrations of: 100μM DL-2-Amino-5-phosphonopentanoic acid (DL-APV, Sigma, St Louis, MO., Cat. No. A5282) or 50μM (D)-2-Amino-5-phosphonopentanoic acid (D-APV, Sigma, St Louis, MO., Cat. No. A8054). To disinhibit (DI) the preBötC network, synaptic inhibition was blocked by bath-application of both 1μM Strychnine (Sigma, Cat. No. S0532) and 50 μM picrotoxin (Tocris, Cat. No. 1128). Norepinephrine (NE) (Sigma, Cat. No. A7256) was bath-applied at a final concentration of 5 μM. Note that while NE increases sigh frequency, the L-APV enantiomer of DL-APV can block sigh-like activity likely via off-target effects on mGlur8 (Lieske and Ramirez 2006). Therefore, since we used DL-APV to block NMDAR in a subset of experiments, we did not further characterize sigh activity in this study and sigh-like bursts were excluded from analyses of preBötC inspiratory bursting activity.

## Data Analyses

### PreBötC Neuronal Activity

The ClampFit 11.2 (Molecular Devices) event detection feature was used to measure the integrated preBötC (∫preBötC) network burst amplitude and frequency. The ∫preBötC network burst amplitude was measured and represented as the difference between the baseline trace value under baseline (ACSF) conditions and the peak height under either control conditions or following drug application. The ∫preBötC network frequency is calculated based on peak ∫preBötC network burst intervals.

To analyze whole cell patch clamp recordings, ClampFit 11.2 (Molecular Devices, San Jose CA, USA) event detection was used to measure action potential frequency during preBötC network bursts and the amplitude and area of inspiratory drive potentials (IDP) and inspiratory drive currents (IDC) that give rise to the depolarization underlying inspiratory neuron bursts. Inspiratory Drive Potentials and Current filtering: To evaluate changes in inspiratory drive potentials (IDP) and currents (IDC) in baseline conditions (ACSF) and following consecutive drug treatments, voltage or current traces were filtered using the ClampFit 11.2, Bessel, 10Hz Low-Pass filters to remove high-frequency transients. Five consecutive inspiratory drive potentials, or currents were averaged during eupneic-like activity and compared between treatments. To maintain temporal alignment of IDC or IDP with ∫pre-BotC network bursts, ∫pre-BotC network bursts activity was also Low-Pass Bessel filtered at 10Hz using ClampFit 11.2.

### Multi-Electrode Array (MEA) data analyses

To analyze MEA data, Tucker-Davis Technologies (TDT) recordings, TDTexport tools were used (see https://tdt.com/docs/sdk/offline-data-analysis/data-conversion/#scaling-notes) to export the data as an interlaced binary file and the data was subsequently imported into the Plexon Inc. (Dallas, Texas, USA) Offline Sorter application (version 4.7.2) via the Plexon software import menu, selecting the binary file with continuously digitized data option. Plexon Offline Sorter was then used to filter the electrical data with a 4-pole Bessler filter (250 Hz low-pass). To sort channel units, we used Plexon Offline Sorter software threshold detection and initially screened units via the Plexon Offline Sorter automated K-means sorting detection function to identify and sort spike events. Subsequent template spike sorting, PCA cluster analysis and visual spike sorting were used to verify channel units. Two minutes of baseline data (in ACSF) was compared to a two minute time course starting 8mins after beginning recordings in NE or APV. To evaluate fidelity of neuron activity, MEA recordings of inspiratory units having at least three (3) or more action potentials during pre-BotC population bursts were selected for analysis. Low fidelity (LF) units were defined as having 3 or more action potentials participating in <95% of preBötC network burst cycles and units were defined as High fidelity (HF) units by having 3 or more action potentials participating in >95% of preBötC network burst cycles. Inspiratory unit percent participation (% participation) was calculated based on the percentage of pre-BotC network bursts (during the last 2mins in each condition - baseline, NE, APV) when an inspiratory unit fired three or more action potentials during each burst.

MEA data were postprocessed in MATLAB (MathWorks, Inc, Natick, MA, U.S.A.). MEA spike times were converted into a population activity signal by binning spikes across all units using 20 ms time bins and normalizing by the number of recorded units, yielding an average network firing rate in unit of spikes/(s · neuron). Network burst onset was detected at time points where the network amplitude crossed a predefined spike threshold from below (default value: 10 spikes/(s · neuron)), and burst termination was defined as the subsequent crossing of this threshold from above. To exclude spurious events, detected bursts were required to exceed a minimum duration of five time bins. For each detected burst, the burst peak was identified as the time point of maximal smoothed population activity within the burst window.

To characterize variability in spike timing around burst events in the MEA data, we constructed ***violin plots*** by aligning neuronal spike times to the burst peak for each detected burst cycle and pooling data across all cycles. For each neuron, spike times within each burst cycle were expressed relative to the corresponding burst peak and normalized by the burst duration. The ***peak action potential density time*** was defined as the time relative to the burst peak at which the spike density reached its maximum across all burst cycles, while the ***mean action potential firing time*** was defined as the average spike times relative to the burst peak.

## Data Analysis Statistics

Statistics were performed using Origin 8 Pro (OriginLab, Northampton, MA, U.S.A.; RRID: SCR_014212) or Prism 6 (GraphPad Software, Boston, MA, U.S.A.; RRID: SCR_015807). In cases where the distribution of data appeared normal, comparisons between two groups were conducted using either paired two-tailed t-tests. A one-way ANOVA was performed followed by Tukey test comparing experimental groups to control for a comparison of three or more groups. Significant differences are reported when the p-value was less than 0.05.

### PreBötC to XII transmission

Transmission from the preBötC to the hypoglossal (XIIn) was expressed as a percentage of the hypoglossal network bursts corresponding to the total network bursts from the preBötC. Bursts were considered corresponding if the initial start time of bursts were within 500–750ms of each other (corresponding time was maximized until only one hypoglossal burst per preBötC was detected). Mean I/O and transmission values for each slice were calculated over a 120 s period taken at the end of each baseline or pharmacological agent phase (each phase duration = 600s). The input-output (I/O) ratio for each inspiratory event (defined by a network burst in preBötC) was calculated as the ratio of preBötC burst amplitude to corresponding hypoglossal burst amplitude (BA) as previously described (Garcia et al., 2016). Calculation of the I/O ratio, was performed using the following equation: IOn = BAXIIn /BA preBötCn, where IO n is the I/O ratio of the n th cycle, BAXIIn is the integrated burst amplitude in the hypoglossal nucleus of the n th cycle and preBötC the integrated burst amplitude in the preBötC of the n th cycle. In cycles where preBötC did not correspond with hypoglossal output, BAXIIn was assigned a value of 0. Prior to the calculation of the I/O ratio, each BAXII was normalized to the mean hypoglossal integrated burst amplitude of the analysis window and each BApreBötCn was normalized to the mean preBötC integrated burst amplitude of the analysis window. Heat maps were used to illustrate individual I/O ratios for up to 25 consecutive cycles in the analysis window for each experiment performed. To illustrate the cycle-to-cycle input-output relationships between networks, heat maps of I/O ratio values were plotted for each slice included in the experiment. Each row represents sequential cycles from a single slice experiment. As the rhythmic frequency across preparations varied, the number of events (i.e. cycle number) in the 120 s analysis window also varied; therefore, either the total number of cycles or up to 25 consecutive cycles from a given slice recording were plotted.

### In Silico Modeling of the preBötC Network

A Hodgkin-Huxley style conductance-based model as previously described in (Jasinski et al., 2013; Phillips and Rubin, 2022) was adopted as the framework for our modeling activities. The currents include a spike generating Na^+^ current (INa), a persistent sodium current (INaP), a delayed rectifying K^+^ current (IK), a voltage-gated Ca^2+^ current (ICa), a Ca^2+^-activated nonspecific cation current (ICAN), a K^+^ dominated leak current (ILeak), a tonic excitatory synaptic current (ITonic) and dynamic excitatory synaptic currents which mediate network interactions. In addition to fast AMPA/kainate synaptic current (InonNMDA), current from NMDA synapse (INMDA) was added to the model, which not only directly depolarizes the neuron but also contributes to calcium influx through NMDAR-mediated calcium transients, with a percentage (PNMDAca) of INMDA carried by Ca^2+^ ions. Networks contain N=100 neurons with 13% connection probability. Full details of the computational model are given in the Supplementary Materials.

As detailed in the model description, there are multiple sources of Ca^2+^ influx from the extracellular space: (1) voltage-gated calcium channel (CaV model), and (2) synaptic inputs, with kainate (CaK model) and NMDA receptors (CaN model) considered as the two possible candidates for synaptically mediated calcium entry. In the model, the parameters 𝑃_𝐼𝐶𝑎_, 𝑃_𝑁𝑀𝐷𝐴𝑐𝑎_ and 𝑃_𝑛𝑜𝑛𝑁𝑀𝐷𝐴𝑐𝑎_ in the calcium equations (see 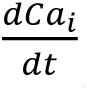 equation in model (1.1) in the Supplementary Materials), were used to control the relative contributions of Ca^2+^ influx via 𝐼_𝐶𝑎_, 𝐼_𝑁𝑀𝐷𝐴_ and 𝐼_𝑛𝑜𝑛𝑁𝑀𝐷𝐴_ currents, respectively. Manipulations of these parameters do not affect these currents in the membrane potential dynamics (see 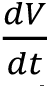 equation in model (1.1) in the Supplementary Materials),. In particular, in the CaV model, calcium influx is generated exclusively from 𝐼_𝐶𝑎_, with 𝑃_𝐼𝐶𝑎_ = 1 while 𝑃_𝑁𝑀𝐷𝐴𝑐𝑎_ and 𝑃_𝑛𝑜𝑛𝑁𝑀𝐷𝐴𝑐𝑎_ set to be zero (i.e., the Ca^2+^ permeability of both NMDA and non-NMDA currents is zero). In the CaK model, (𝑃_𝑛𝑜𝑛𝑁𝑀𝐷𝐴𝑐𝑎_ = 1, 𝑃_𝐼𝐶𝑎_=𝑃_𝑁𝑀𝐷𝐴𝑐𝑎_ = 0), while in the CaN model, (𝑃_𝑁𝑀𝐷𝐴𝑐𝑎_ = 1, 𝑃_𝐼𝐶𝑎_=𝑃_𝑛𝑜𝑛𝑁𝑀𝐷𝐴𝑐𝑎_ = 0).

Heterogeneity was introduced by normally distributing the parameters 𝑔_𝐿_ and 𝑔_𝑁𝑎𝑃_. Additionally, 𝑔_𝐶𝐴𝑁_ was uniformly distributed over the range 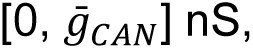 and the weights of excitatory synaptic connections were uniformly distributed as 𝑊_𝑗,𝑖_ = 𝑈(0, 𝑊_𝑀𝑎𝑥_), with 𝑊_𝑀𝑎𝑥_ = 0.15 𝑛𝑆 for non-NMDA receptors and 𝑊_𝑀𝑎𝑥_ = 0.05 𝑛𝑆 for NMDA receptors (see the Supplementary Materials for modeling details). To investigate the effects of NMDA and CAN currents on network dynamics, we varied the maximum CAN conductance and the maximum NMDA synaptic weight across all neurons over the range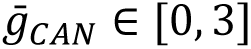 nS and 𝑊_𝑀𝑎𝑥,𝑁𝑀𝐷𝐴_ ∈ [0, 0.1] nS (see Figures 4, 5, 7). In particular, to mimic pharmacological NMDAR blockade experiments, simulations were initialized at an elevated NMDA state (𝑊_𝑀𝑎𝑥,𝑁𝑀𝐷𝐴_ = 0.1 nS) and NMDAR weights were progressively reduced to zero while tracking the resulting changes in network dynamics.

### In Silico Model Data Analysis and Integration methods

Data generated from simulations was postprocessed in MATLAB (MathWorks, Inc). A spike was defined as occurring in a neuron when its membrane potential (𝑉) depolarized passed a threshold of-35 mV. Consistent with the MEA data analysis, the network amplitude, a measure of the average firing rate of the neuronal population, was calculated as the total number of spikes generated by the network per 20 ms bin divided by the number of neurons in the network, with units of spikes/(s · neuron). Network burst amplitudes and frequencies were calculated by identifying the peaks and the inverse of the interpeak interval from the network amplitudes. Simulation software was custom written in C++. Numerical integration was performed using the exponential Euler method with a fixed step-size (Δt) of 0.025 ms.

### Synchrony Score

Spike rasters (obtained from either model simulations or from MEA recordings) were stored as a sparse matrix, indicating whether or not each neuron fires a spike at a given time step. To reduce its size, the spike raster data are aggregated into 50-ms bins of spike counts. This produces a vector time series 𝑋(𝑡) = (𝑥_1_(𝑡),…, 𝑥_𝑁_(𝑡)), where 𝑥_𝑖_(𝑡) represents the spike count of the 𝑖-th neuron at bin time 𝑡. To characterize the overall synchrony of the population, we used the following statistic, adapted from Golomb 2007, Masuda and Aihara 2004 and Harris et al. 2017:

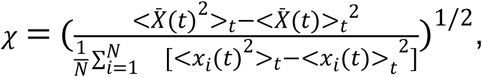

where the angle brackets <. > denotes averaging over the time series and 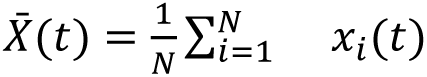 denotes averaging across all neurons in the population.

### Experimental data and code accessibility

The simulation code, experimental data and data processing code will be made available upon reasonable request.

## Results

### Norepinephrine modulation increases preBötC burst amplitude and frequency

To investigate the role of norepinephrine (NE) modulation of the preBötC network, we made single electrode extracellular recordings of rhythmic inspiratory bursts generated by the preBötC network isolated in medullary brainstem slice preparations; recordings of inspiratory preBötC network bursts were made prior to (**Fig 1A, Baseline**) and following bath application of 5µM NE (**Fig 1B, NE**). Consistent with prior reports, (Viemari and Ramirez 2006; Zanella et al. 2014), NE increases both integrated preBötC network burst amplitude (**Fig. 1C**) and preBötC network burst frequency (**Fig. 1D**).

**Figure 1.**
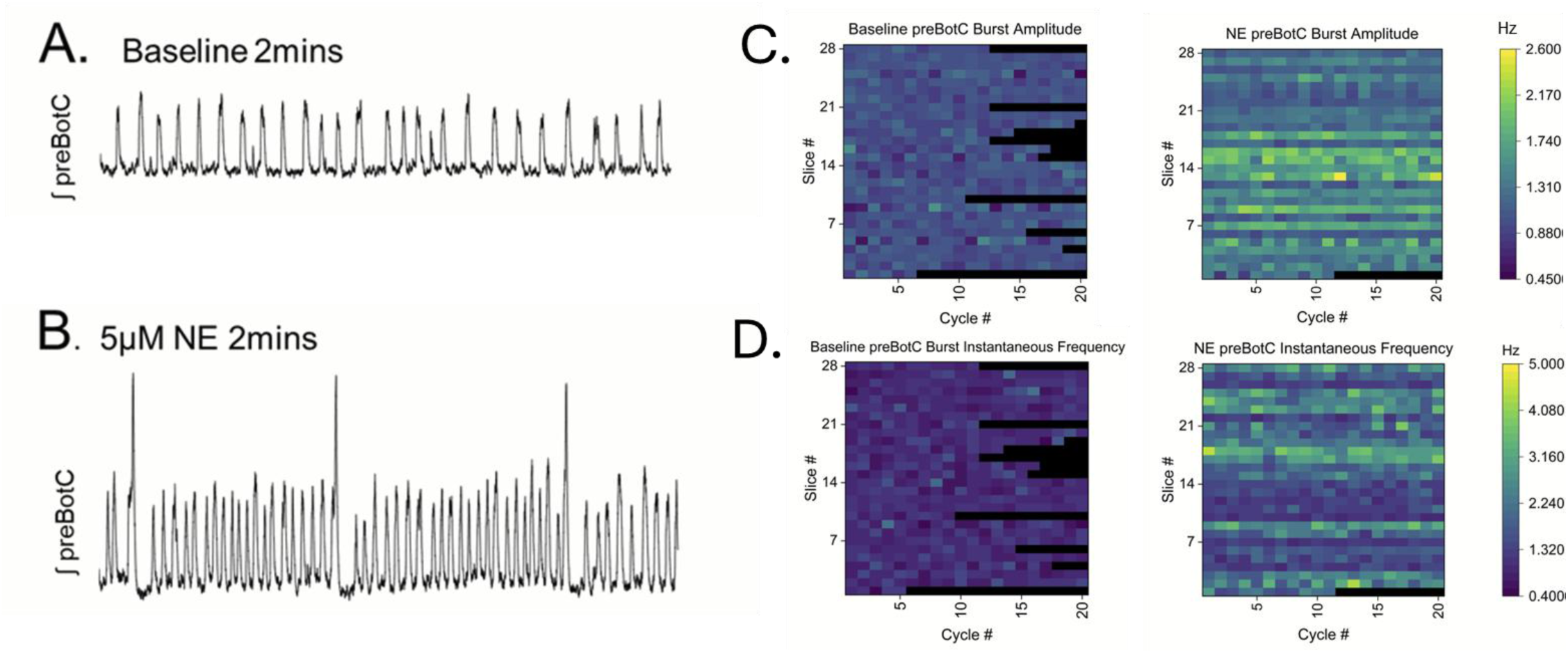
Norepinephrine (NE) modulation enhances pre-Botzinger Complex (preBötC) network burst amplitude and burst frequency. **(A)** Integrated preBötC network bursting under baseline conditions. **(B)** preBötC network burst amplitude and frequency increases following bath-application of 5µM NE. As in prior studies (Viemari and Ramirez 2006), large amplitude fictive sigh burst frequency increases with NE. Note that sigh-like activity was excluded from analysis (see Methods). **(C)** Heatmaps show that compared to baseline conditions (left heatmap), preBötC network burst amplitude increases during NE modulation (right heatmap) (preBötC burst amplitude in baseline vs. NE, p=0.00173, paired T-test, average NE amplitude = 139.904% ±26.808 SD above baseline (100%), n=28 slices); and, **(D)** pre-BotC instantaneous frequency increases with NE modulation (p<0.0001, paired T-test; mean IF baseline = 0.0239 ± 0.136 SD vs mean NE 0.458 ± 0.221 SD; n=28 slices). Black color indicates the preBötC network did not generate a burst during a cycle. Heatmaps and analysis include n=28 slice preparations (Slice # 1-28, y-axis) and included up to 20 consecutive preBötC network bursts from each slice (Cycle #1-20, x-axis).

### NE modulation increases preBötC burst amplitude via enhanced NMDAR activity

Following NE modulation, the enhanced preBötC network burst amplitude suggested the possibility that NE enhances inspiratory network synchronization. Given the recognized role of NMDAR in mediating neural network synchrony throughout the brain (Flint et al. 1996; Takahashi et al. 2012; Gu et al., 2013), we tested whether blockade of NMDAR with the NMDAR antagonist APV impacts the preBötC eupneic-like rhythm following NE modulation (n=12 slices). NE modulation increases both preBötC burst amplitude and frequency (**Figs. 2A-2C**), subsequent blockade of NMDAR with APV activity reduces burst amplitude (**Fig. 2B** right). Paradoxically, blocking excitatory glutamate NMDARs with APV increases preBötC network burst frequency in the majority of slices (n=10/12; 83.3%) and preBötC network burst frequency remains elevated above baseline frequency (**Figs. 2B-2C**).

**Figure 2.**
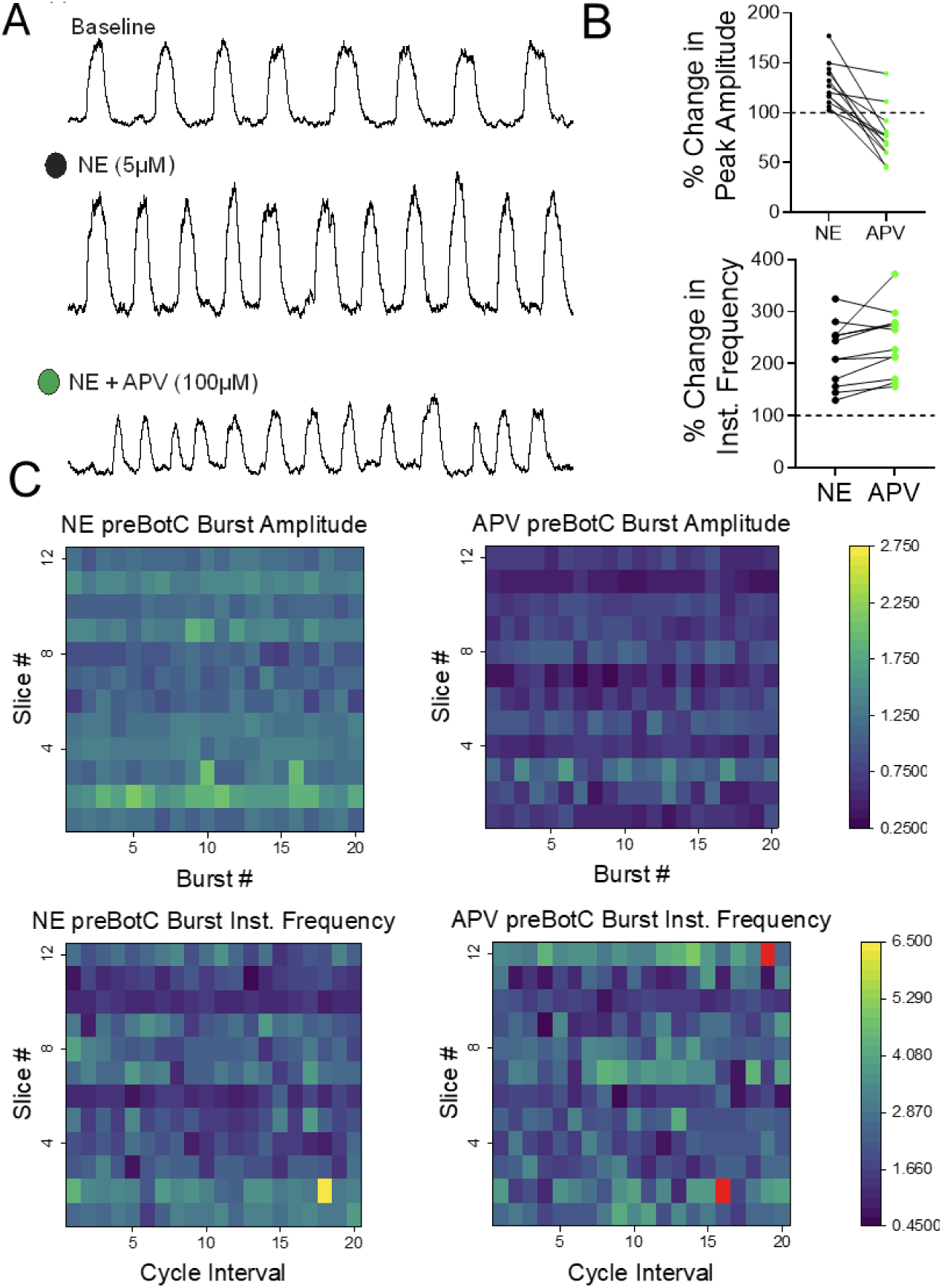
Norepinephrine-mediated signaling enhances burst amplitude which is subsequently reduced by blocking NMDAR. **(A)** Integrated preBötC network bursts under baseline conditions (top trace), after bath-application of 5µM NE (middle trace) and following blockade of NMDAR (bottom trace); 30s of activity are shown in each condition. **(B)** PreBötC burst amplitude and frequency increases above baseline values following bath-application of 5µM NE. PreBötC burst amplitude increased to 128.55% ± 21.42 mean ± SD, p=0.00302 relative to baseline (100%) RM ANOVA, Tukey, n=12 slices) and burst frequency increased to 118.97% ± 59.76 mean ± SD, p=<0.0001, RM ANOVA, Tukey n=12 slices). Blocking NMDAR with APV subsequently reduces burst amplitude to 77.54% ± 26.79526 mean ± SD relative to baseline (100%), p=0.0193, RM ANOVA, Tukey, and by 51.00% ± 30.07 mean ± SD relative to NE, p=<0.0001 one-way RM ANOVA, Tukey, n=12 slices). During NE modulation, blocking NMDAR led to an increase in preBötC burst instantaneous frequency (IF) in the majority of slices (n=10/12; 83.33%). In n=2/12 slices (16.66%), preBötC IF decreased after blocking NMDAR; yet preBötC burst mean IF remained elevated above baseline levels in all slices (n=12/12; p<0.0001; at 142.41% ± 63.77 mean ± SD relative to baseline, RM ANOVA, Tukey). **(C)** Heatmaps illustrate that compared to baseline conditions (left heatmaps) both burst amplitude (top row) and burst frequency (bottom row) increase with NE mediated signaling. Subsequent blockade of NMDAR with APV reduces burst amplitude and typically enhances burst frequency (RED color indicates value is >6.5). Heatmaps include n=12 slice preparations (slices #1-12, y-axis) and up to 20 consecutive bursts (burst #1-20, x-axis). Quantitative graphs compare burst amplitude and frequency during 2mins of steady-state baseline, NE or APV conditions in n=12 slices.

Following NE modulation, we considered the possibility that after blocking NMDAR, the significantly decreased preBötC network burst amplitude, could result in failure of the preBötC network to activate downstream motor pools, such as the hypoglossal nucleus (XIIn) which plays a critical role in upper airway patency. Indeed, small amplitude preBötC network bursts have been associated with the occurrence of subnetwork activity whereby the preBötC network fails to evoke output at the level of the hypoglossal nucleus (XIIn) (Zanella et al. 2014; Browe et al. 2023; Garcia et al. 2016; Kallurkar; et al. 2020). Therefore, we performed simultaneous network recordings from the preBötC and XIIn to determine how blockade of NMDAR impacted the efficacy of the preBötC to drive XII motor output (**Supplemental Fig 1A**; n=3). While APV reduced preBötC network burst amplitude (**Supplemental Fig. 1B**), the one-to-one relationship between the preBötC network burst and XIIn output was preserved and the input-output ratio between the preBötC and XIIn was unchanged (**Supplemental Fig. 1C**), indicating that while following NE modulation, subsequent loss of NMDAR activity produces small amplitude preBötC network bursting, it does not lead to subnetwork activity that fails to generate downstream XII activity.

To determine whether synaptic inhibition of the preBötC is preserved following NE-modulation, we performed a series of experiments where we disinhibited the network following NE-modulation (**Fig. 3A**). Disinhibition (DI) stimulated burst amplitude (**Fig. 3B**) while reducing burst frequency of the preBötC network rhythm (**Fig. 3C**). These observations suggest that intact synaptic inhibition contributes to shaping the network response to NE. Given this possibility, we sought to determine whether NMDAR recruitment is an indirect phenomenon caused by a shift in E/I balance, we conducted a series of whole-cell patch clamp recordings (n=9 total, with n=5 in voltage clamp configuration and n=4 in current clamp configuration) where inspiratory drive potentials (IDP; **Fig. 4A-4D**) and inspiratory drive currents (IDC; **Fig 4E-4G**) were measured. Relative to baseline conditions (**Fig 4A-C, red**), DI increased the area of the IDP (**Fig 4A-C, blue**) but on average did not affect the number of action potentials during the preBötC network burst **(Fig. 4D)**. Consistent an increase in IDP area following DI, DI also enhanced the IDC area (**Fig. 4E-4G**, **red vs. blue**). Subsequent NE modulation increased IDP area (**Figs 4A-C**, **black**) and this enhanced excitability increased the number of action potentials generated during preBötC network bursts relative to baseline conditions **(Fig. 4D)**; consistent with this enhanced excitability, NE modulation also enhanced the IDC area and IDC duration (**Figs. 4E-4G**, **black**). Although following DI, NE increased the mean number of action potentials per burst above baseline conditions, NE increased action potential firing above DI levels in n=3 of 4 neurons, while one neuron had a lower number of action potentials following NE modulation, versus in the DI state alone (**Fig. 4D**); these data suggest there may be a heterogeneity in individual neuron activity in response to NE modulation.

**Figure 3.**
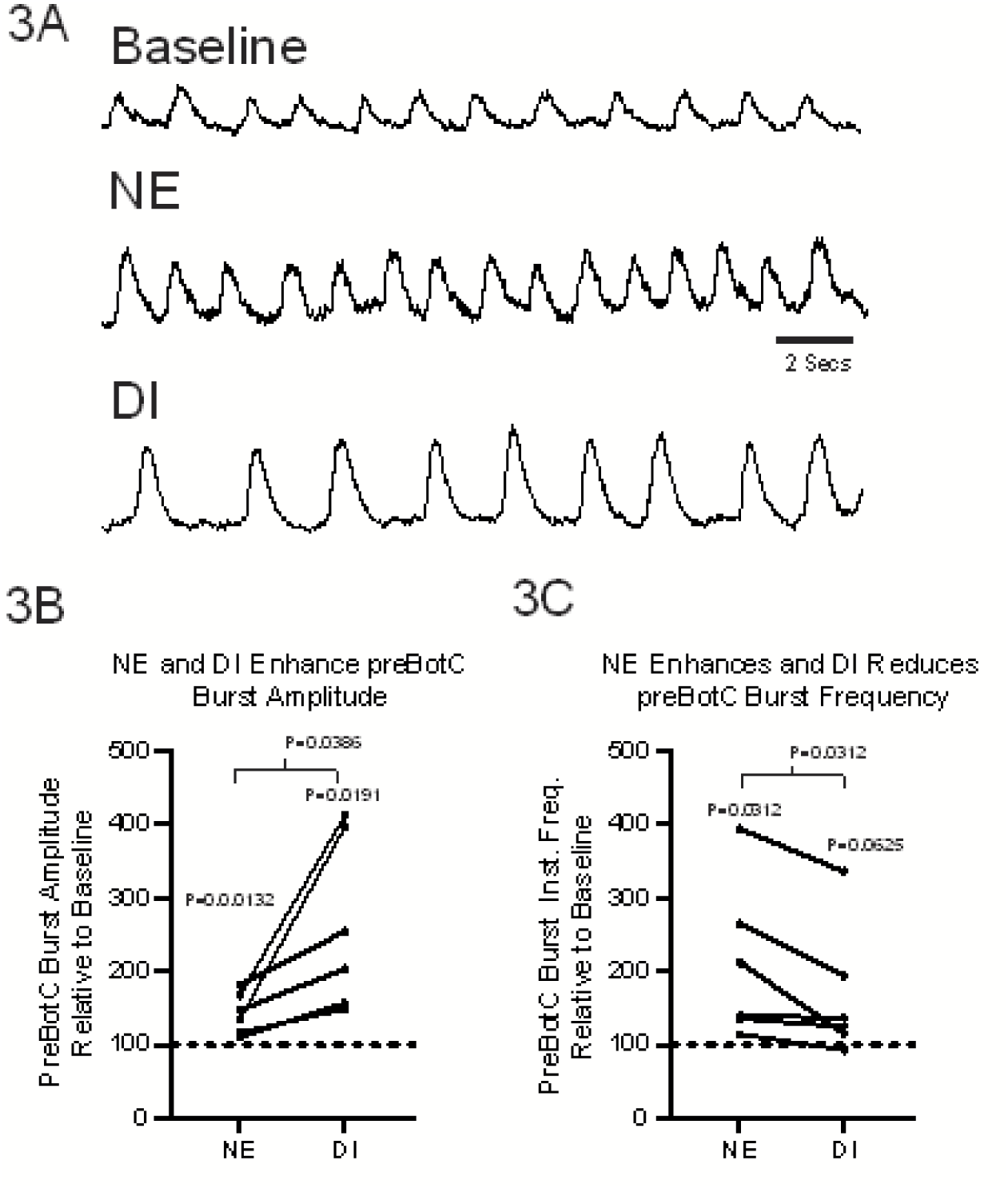
Synaptic inhibition remains intact following norepinephrine (NE) modulation. **(A)** Compared to baseline (acsf), following bath-application of 5µM NE, integrated preBötC network burst **(B)** amplitude and **(C)** frequency increase above baseline. In NE PreBötC burst amplitude increased above baseline by 43.06% ± 24.47 mean ± SD, p=0.0132 paired t-test). In NE, PreBötC burst frequency increased by 109.90% ± 106.097 mean ± SD p=0.312 t-test above baseline, **(A-C).** During NE-modulation, subsequent disinhibition (DI) of the preBötC results in a further increase in burst amplitude (by 81.23% ± 24.4 mean ± SD above baseline, p=0.0191, RM ANOVA, Tukey) but (Fig 3C) a reduction in burst frequency (by-66.16% ± 89.29 mean ± SD, p=0.0017, RM ANOVA, Tukey).

**Figure 4.**
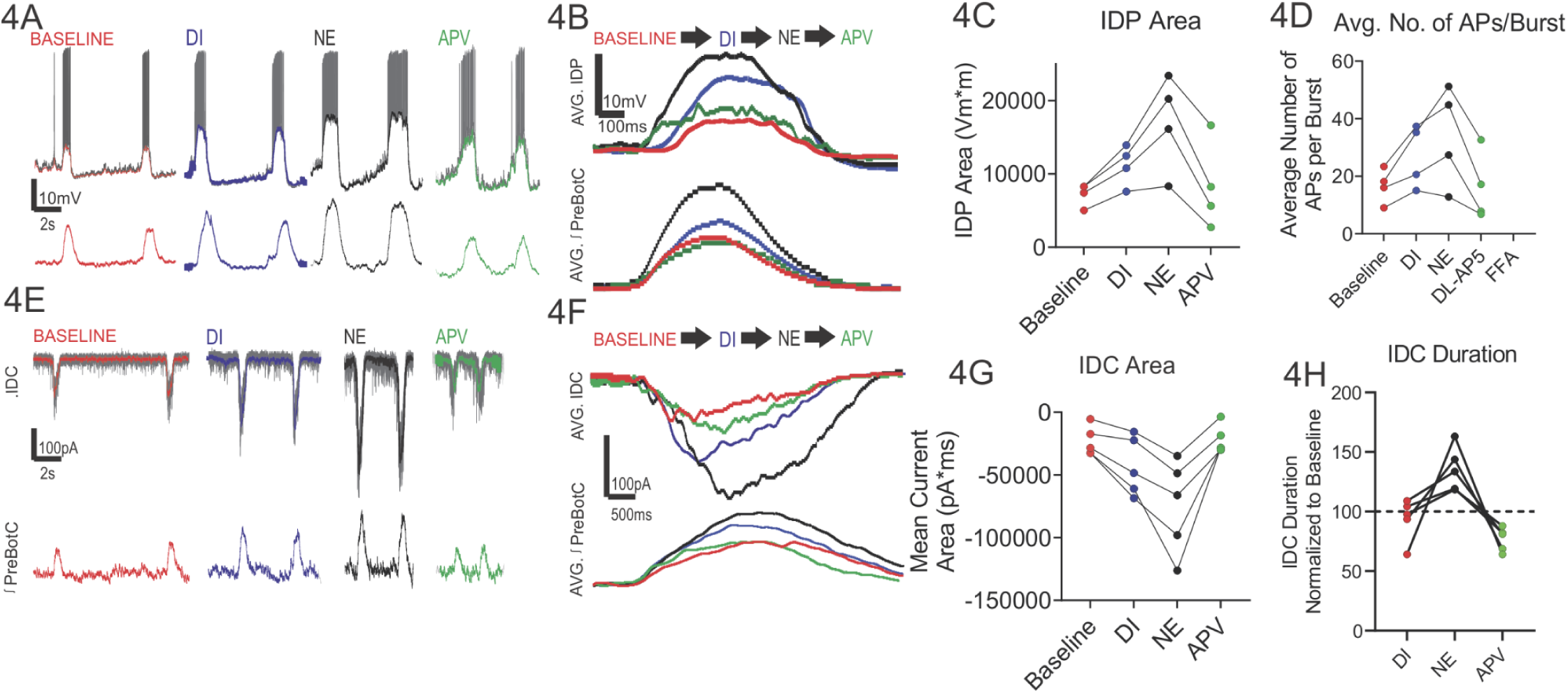
N**o**repinephrine **(NE) signaling enhances preBötC inspiratory drive potentials (IDP) and currents (IDC) and neuron excitability via NMDAR activity. (A)** Current clamp recording (top row) of inspiratory neuron potentials and preBötC population bursts (bottom row) under baseline conditions, following disinhibition (DI, blue), norepinephrine (NE) modulation (black) and following blockade of NMDAR (APV, green). The (filtered) IDP underlying each burst, under each condition is overlaid on top of the current clamp trace. **(B)** Averaged IDP and population bursts reveal **(C)** the IDP area increased following DI (p=0.00958, paired T-test; mean Baseline = 7258 Vm*ms ± 1537 SD vs DI mean = 11191 Vm*ms ± 2726 SD) but **(D)** the number of spikes per burst did not significantly increase following DI (p=0.1706, One way RM ANOVA; mean Baseline = 16.65 ± 5.97 SD; mean DI=27.05 ± 10.96 SD). Subsequent NE modulation increased both (Fig. 4C) the IDP area (baseline vs. NE, p=0.0031, one-way RM ANOVA; NE mean = 17029 Vm*ms ± 6517 SD) and (Fig. 4D) average number of action potentials per burst with respect to baseline conditions (APs/burst, baseline vs. NE, p=0.0171, one-way RM ANOVA; NE mean = 34.05 ± 17.37SD). Fig. 4C, Blockade of NMDAR with APV decreased both the IDP area (NE vs. APV, p=0.00649; APV vs baseline, p=0.94588; one way RM ANOVA; APV mean = 8308 mV*ms ± 5989 SD); and, **(D)** average number of action potentials per burst (NE vs APV, p=0.01434; APV vs. Baseline, p=0.999; one way RM ANOVA; APV mean = 16.10 +/-11.96 SD). **(E)** Voltage clamp recordings (top row) of inspiratory neuron currents and preBötC population bursts (bottom row) under baseline conditions, following disinhibition (DI, blue), norepinephrine (NE, black) modulation and following blockade of NMDAR (APV, green). The (filtered) IDC underlying each burst under each condition is overlaid on top of the voltage clamp trace. **(F)** Averaged IDC and population bursts reveal **(G)** the IDC area increases from baseline following DI (p=0.025, baseline vs. DI, paired T-test, n=5), increases further with NE modulation (p=0.026, DI vs NE, RP ANOVA, n=5) and is reduced to baseline levels following blockade of NMDAR with APV (p=0.996, baseline vs. APV RP ANOVA, n=5). **(H)** The duration of the IDC does not change following DI (duration in DI was 93.7% ±17.55 SD of baseline, p=0.603 One-way RM ANOVA, n=5) but increases following NE modulation (duration in NE was 35.6% ± 18.61 SD of baseline, One-way RM ANOVA, n=5) and is not significantly different than the baseline duration following blockade of NMDAR with APV(duration in APV was 76.80% ± 9.962 SD of baseline, p=0.86565, One-way RM ANOVA, n=5).

Our preBötC network recordings suggested NE enhances preBötC network bursting via increasing NMDAR activity. To test if NE modulation recruits NMDAR activity to enhance inspiratory neuron excitability, we next blocked NMDAR with APV which reduced both the IDP (**Fig. 4A-4G, green**) and IDC (**Fig. 4E-4G**) and reduced the number of action potentials fired during preBötC network bursts (**Fig. 4D**) and reduced IDC duration **(Fig. 4H)**. These data demonstrated that NE modulation of inspiratory neuron activity via NMDAR recruitment can play a critical role in modulating synaptic activity within the preBötC and NMDAR recruitment can be evoked even in states where inhibition is blocked. These observations also raise the possibility that synaptic inhibition suppresses recruitment of NMDAR as (after NE modulation) subsequently blocking NMDAR reversed the effects of DI to near baseline values **Figs. 4A-4H**.

### In silico experiments predict NE-modulation of NMDAR effects on network behavior

Synaptic sources of intracellular calcium ([Ca^2+^]_i_) have been proposed as a critical regulator of network activity (Philips et al. 2019). In a series of in silico experiments, we next examined how distinct modes of Ca^2+^ entry influenced network responses to NMDAR blockade initiated from an NE-like state. We considered three network configurations: (i) a CaV model in which calcium influx occurred exclusively through voltage-gated Ca^2+^ channels; (ii) a CaK model where synaptic calcium influx occurred exclusively through Ca^2+^-conducting non-NMDAR glutamate receptors; and finally, (iii) a CaN model in which NMDAR-mediated calcium influx served as the sole synaptic source of elevated [Ca^2+^]_i_. In the CaV model network, the impact of 𝑊_𝑀𝑎𝑥,𝑁𝑀𝐷𝐴_on network dynamics arose solely from NMDAR-mediated membrane potential depolarization, since NMDARs were made not Ca^2+^-permeable. With gCAN fixed 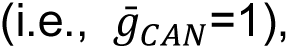 decreasing the weight of the NMDAR across the network modestly decreases network burst frequency (Fig. **5A** top row, red curve). This frequency decrease resulted from a slight reduction in intracellular calcium concentrations (**Fig. 5A**, third row), caused by diminished NMDAR-mediated depolarization. The lower calcium levels, in turn, decreased the amplitude of ICAN during the interburst interval (**Fig. 5A** bottom row), making neurons less able to reach burst threshold each cycle and thereby contributing to a lower burst frequency. On the other hand, modulating NMDAR weight had negligible influence on network burst amplitude (**Fig 5A** top row, blue) or on spike timing of individual neurons as illustrated in the corresponding raster plot (**Fig. 5A** second row). These findings are consistent with previously reported effects of ICAN on a CaV network (Phillips et al., 2019), though the changes we observe here are much smaller because ICAN is affected indirectly only through NMDAR-mediated voltage changes in these networks. In the CaK model network, where the source for the rise of [Ca^2+^]_i_ was solely derived from Ca^2+^-conducting non-NMDAR glutamate receptors, such as kainate receptors (Paarmann et al., 2000; Perrais et al., 2009, Philips et al. 2019), varying 𝑊_𝑀𝑎𝑥,𝑁𝑀𝐷𝐴_ had negligible effects on network dynamics (**Fig 5B**). These outcomes illustrate the insensitivity of NMDAR modulation on the dynamics of both CaV and CaN networks, an effect that persisted across a range of 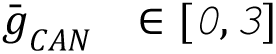 (see **Supplementary Figure 2**), and similarly held when [Ca^2+^]_i_ elevations were driven jointly by voltage-gated and non-NMDA synaptic sources, regardless of their relative contributions (*data not shown*). Together, these observations suggest that non-NMDAR dependent sources of elevated [Ca^2+^]i are not sufficient for driving the network dynamics experimentally observed with NE (**Fig. 2**).

**Figure 5.**
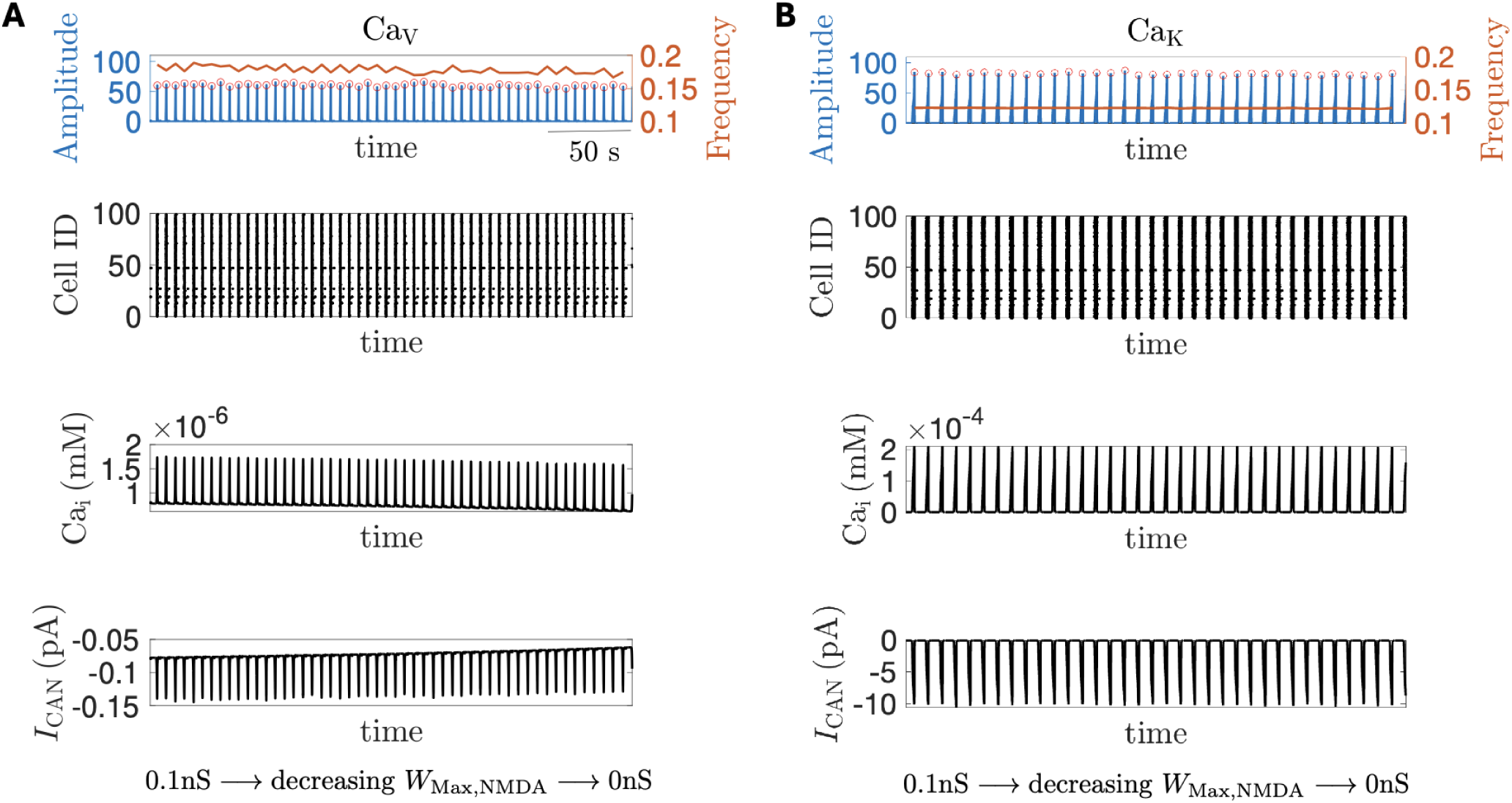
When NMDA receptors are impermeable to Ca^2+^, varying NMDAR coupling strength has minimal impact on network activity. In both **(A)** CaV and **(B)** CaK networks, linearly varying NMDAR weight 𝑊_𝑀𝑎𝑥,𝑁𝑀𝐷𝐴_ had minimal effects on (first row) neuronal population activity amplitude (blue curve) and frequency (red curve); (second row) average intracellular calcium concentration (𝐶𝑎_𝑖_); and (bottom row) average 𝐼_𝐶𝐴𝑁_ current.

In the CaN model network, where NMDAR are permeable to Ca2+, a greater impact of NMDAR coupling strength on network dynamics was observed. Consistent with experimental observations (**Fig. 2**), decreasing 𝑊_𝑀𝑎𝑥,𝑁𝑀𝐷𝐴_ increased network burst frequency (**Fig. 6A** top row) while reducing network burst amplitude (**Fig 6A** bottom row) and network synchrony (**Fig. 6B**). Since in our model, [Ca^2+^]_i_ elevations influence network dynamics by activating ICAN, reducing 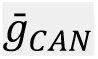 attenuates the influence that 𝑊_𝑀𝑎𝑥,𝑁𝑀𝐷𝐴_ exerts on the network.

**Figure 6.**
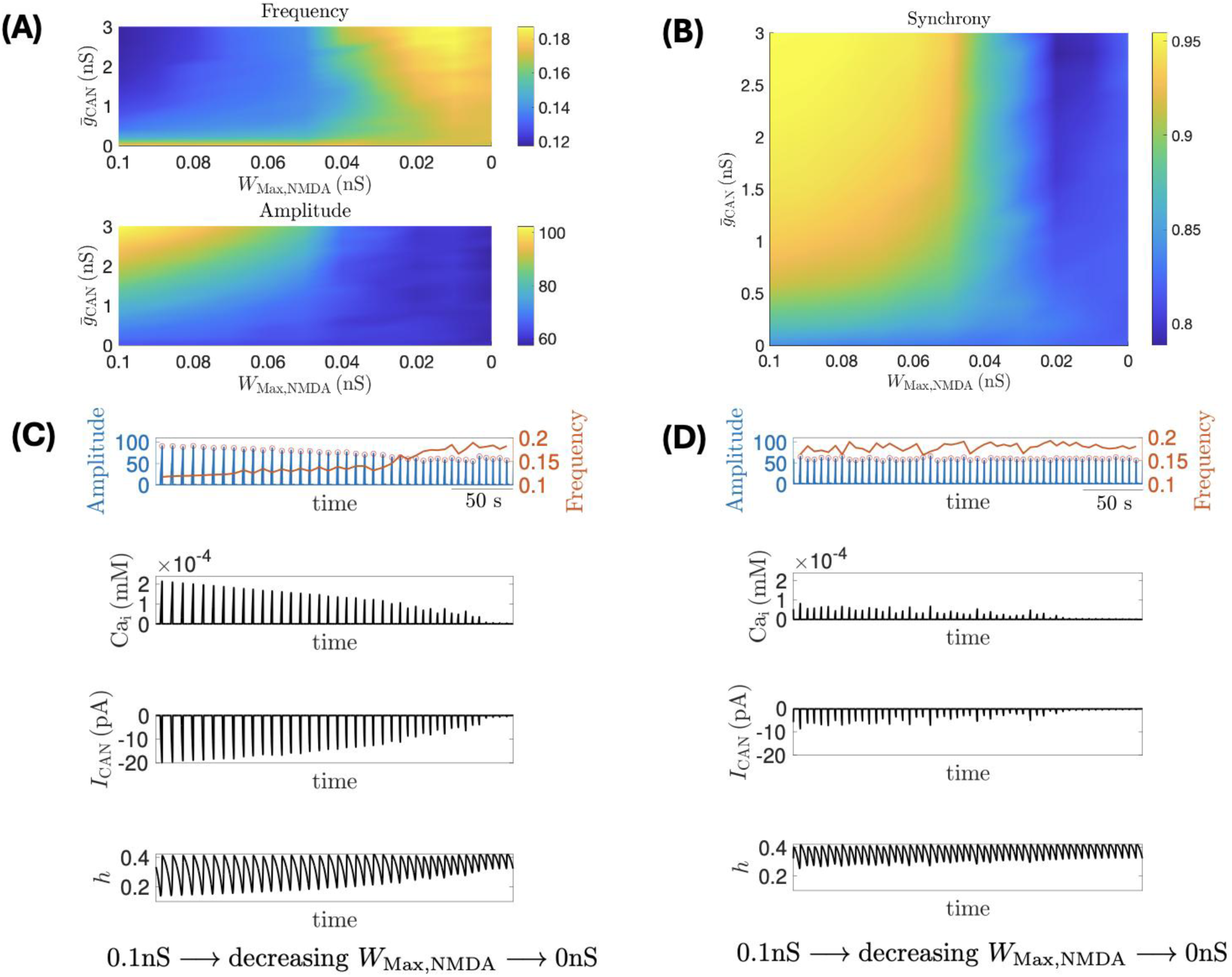
Effect of varying NMDAR weight on the CaN network, where calcium influx is exclusively mediated by NMDAR synaptic input. (A; **B)** Decreasing 𝑊_𝑀𝑎𝑥,𝑁𝑀𝐷𝐴_ in the CaN network increases burst frequency while decreasing network amplitude and synchrony. These effects are mitigated with reduced 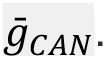 **(C; D)** Each panel consist of: (top row) histograms of neuronal population activity amplitude (blue) and frequency (red) at 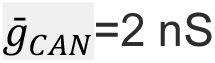 with linearly decreasing 𝑊_𝑀𝑎𝑥,𝑁𝑀𝐷𝐴_; (second row) average intracellular calcium concentration (𝐶𝑎_𝑖_); (third row) average 𝐼_𝐶𝐴𝑁_ current; (bottom row) average inactivation (ℎ_𝑁𝑎𝑃_) of the burst generating current 𝐼_𝑁𝑎𝑃_. **(C)** NMDAR decay constant 𝜏_𝑁𝑀𝐷𝐴_ = *`20`* ms; **(D)** 𝜏_𝑁𝑀𝐷𝐴_ = *`5`* ms. Other parameters are given in Methods.

To gain a better understanding of the somewhat counterintuitive increase in network frequency as the NMDAR synaptic weight is reduced, we fixed 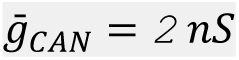 and examined the effect of linearly decreasing 𝑊_𝑀𝑎𝑥,𝑁𝑀𝐷𝐴_ on network amplitude frequency (**Fig 6C top row**), average intracellular calcium concentration (𝐶𝑎_𝑖_, **Fig 6C, second row)**, CAN current (𝐼_𝐶𝐴𝑁_, **Fig 6C, third row**) and inactivation kinetics of the NaP current (ℎ, **Fig 6, bottom row**), which can largely determine burst network frequency (Philips et al. 2019, Wang and Rubin 2016). Decreasing 𝑊_𝑀𝑎𝑥,𝑁𝑀𝐷𝐴_ reduces [Ca^2+^]_i_ during bursts (**Fig. 6C, second row**), which in turn lowers ICAN activation (**Fig. 6C, third row**). The reduced depolarization within each burst naturally leads to smaller network amplitudes and also disrupts synchrony across neurons, consistent with the effects of reducing gCAN which directly decreases ICAN.

The more interesting effect concerns network burst frequency. Reduced depolarization within bursts raises the hNaP value at which bursts terminate, as indicated by the progressively higher trough ℎ-values in the bottom row of **Fig. 6C**. In contrast, 𝑊_𝑀𝑎𝑥,𝑁𝑀𝐷𝐴_ has little effect on 𝐶𝑎_𝑖_ and ICAN between bursts (**Fig. 6C**, third row), leaving the hNaP threshold for the burst initiation unaffected, as indicated by the nearly constant peak values of ℎ in **Fig. 6C** bottom row. Because the initiation threshold remains nearly constant, a higher termination threshold allows h to reach it sooner as it decreases during the burst, leading to earlier termination and earlier onset of the next burst. This shortens the interburst interval and increases the network burst frequency as 𝑊_𝑀𝑎𝑥,𝑁𝑀𝐷𝐴_ decreases.

The effects mediated by NMDAR above are primarily due to the relatively slow decay of NMDAR currents (∼20 ms; Paarmann et al., 2005; Morgado-Valle and Feldman, 2007). When the decay time is shortened to match the fast decay of AMPAR/KAR currents (∼5 ms), 𝑊_𝑀𝑎𝑥,𝑁𝑀𝐷𝐴_ has almost no effect on the network dynamics despite the presence of NMDAR-mediated calcium influx (**Fig. 6D**). The inability for 𝑊_𝑀𝑎𝑥,𝑁𝑀𝐷𝐴_ to reliably influence network dynamics coincided with the limited changes in [Ca^2+^]i, ICAN and hNaP oscillations (**Fig. 6D**), relative to the respective metrics from the CaN model network with a larger NMDAR decay constant (**Fig. 6C**). These findings underscore the importance for an interplay between slower kinetics of the NMDAR currents and [Ca^2+^]i sequestration having a key role in mediating changes to network activity when NMDAR activity is recruited.

### Impact of NE modulation on interneuronal dynamics in the preBötC

To further resolve the impact of NE on the preBötC we performed a series of multi-electrode array (MEA) recordings (n=7 preparations) where the behavior of individual units in the preBötC could be resolved under baseline (**Fig 7A**), during NE (**Fig 7B**) and following subsequent blockade of NMDAR activity (**Fig. 7C**). Consistent with our intracellular observations, NE has a heterogeneous impact on neuronal excitability. While NE modulation reduced the average action potential number during the network burst (**Fig. 7D**), it increased action potential frequency during inspiratory network bursts (**Fig. 7E**). APV application further reduced both action potential number (**Fig 7D**) and suppressed the NE-driven increase in action potential frequency (**Fig 7E**). Relative to Baseline (**Fig 8A**), NE (**Fig 8B-8E**) shifted both mean spike timing and peak action potential density closer to, or ahead of the peak of the network burst. APV (**Figs. 8C-8E**) did not further impact on either spike timing (**Fig 8D**, green) or peak action potential density (**Fig 8E, green**) suggesting that NE-mediated shifts in the timing of action potential firing and elevated action potential frequency during preBötC network bursts persist despite the loss of NMDAR activity.

**Figure 7.**
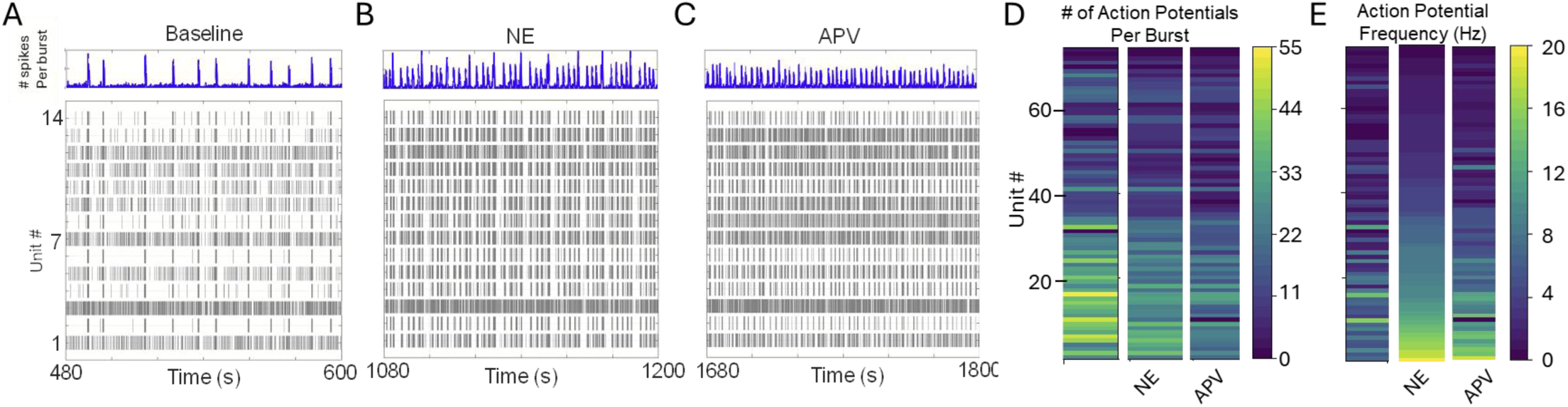
Norepinephrine (NE) modulation and subsequent NMDAR blockade changes preBotzinger (preBötC) inspiratory unit dynamics. Compared to **(A)** baseline experimental conditions, **(B)** NE modulation enhances preBötC network burst amplitude (ʃpreBötC) and frequency, and **(C)** subsequent blockade of NMDA (via APV) reduces network burst amplitude (ʃpreBötC), while burst frequency remains elevated. See also Fig. 2. **(D)** Relative to baseline, both NE modulation and subsequent blockade of NMDAR reduced the number of action potentials per preBötC network burst. Baseline mean 21.474 ± 1.604 SE vs NE mean 16.856 ± 1.205 SE, Baseline vs NE p=<0.001; APV mean = 13.006+/-1.025 SE, NE vs APV, p=<0.0001; APV vs. Baseline p=<0.0001; n=74 units; RM ANOVA, Tukey **(E)** Relative to baseline, NE modulation increases the action potential frequency during preBötC network bursts (p<0.0001; Baseline mean 3.78 (Hz) ± 0.394SE vs. NE mean 6.29 (Hz) ±0.58 SE) and subsequent blockade of NMDAR reduced the average action potential frequency during preBötC network bursts (p=0.00602; mean NE =6.29 (Hz) ±0.58 SE vs mean APV 5.085 (Hz) ±0.521 SE) although in APV, action potential frequency remained elevated above baseline (p=0.00284, APV vs Baseline; RM ANOVA, Tukey, n=74 units).

**Figure 8.**
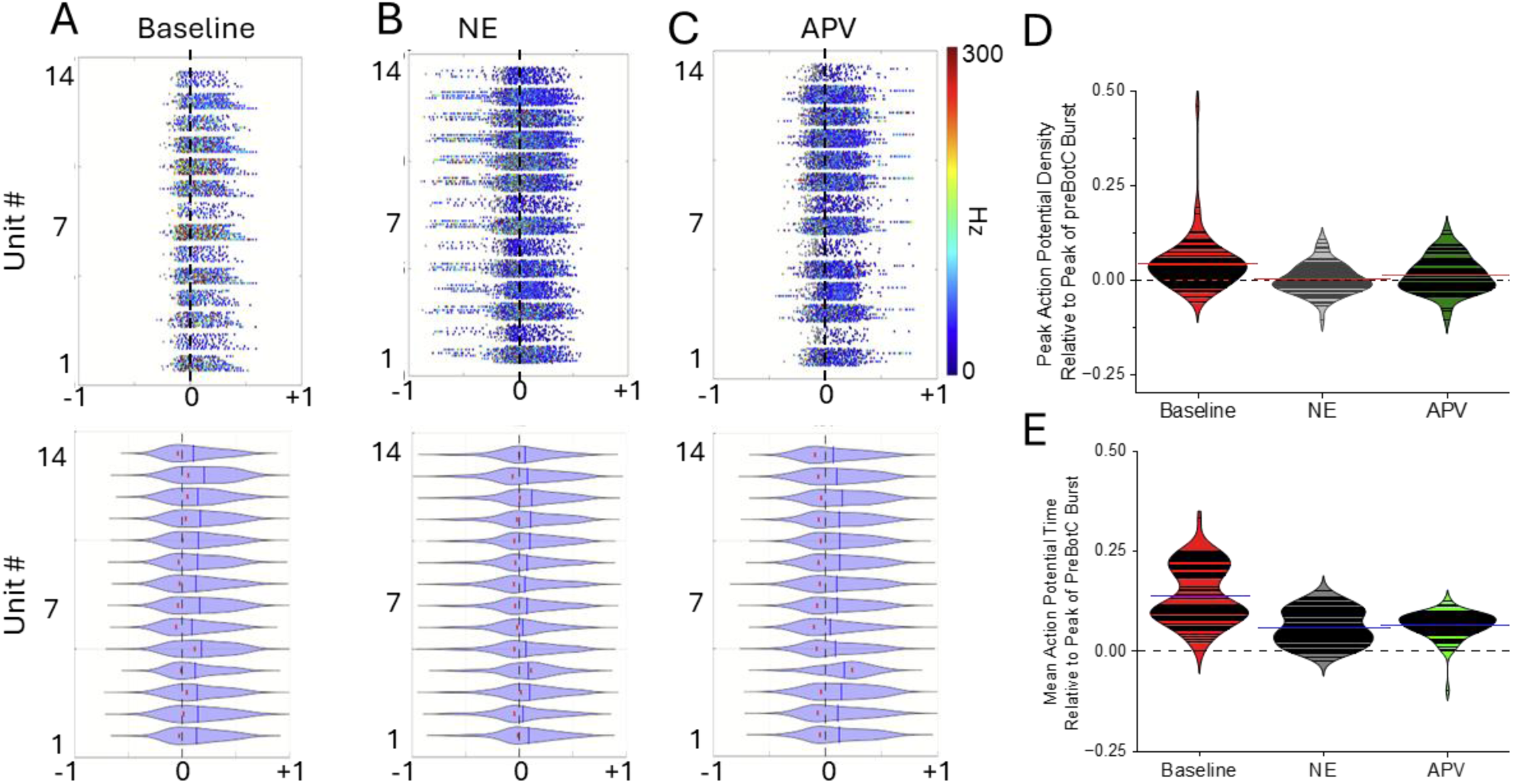
**Norepinephrine (NE) modulation of preBötC inspiratory units shifts their action potential firing pattern toward or ahead of the peak of the preBötC network bursts**. Representative frequency coded raster plots (top) and violin plots (below) of inspiratory unit action potential timing relative to the peak of the preBötC network during **(A)** baseline; **(B)** NE modulation and **(C)** following blocking NMDAR with APV (peak is at x=0, black dashed vertical line). Violin plots reveal peak action potential density (red dots) and mean action potential firing time shifts closer to or ahead of the preBötC network burst peak (x=0)**. (D)** Relative to Baseline (red) peak firing density shifts closer to the peak of the preBötC network burst (at y=0, horizontal black dashed line) during NE modulation (gray) and persists following blockade of NMDAR with APV (green). Peak firing density: Baseline=0.357 ± 0.00565 SE; NE= 0.00289 ± 0.00469 SE; APV= 0.128 ± 0.00564 SE; Baseline vs NE p=<0.001; NE vs.APV p=0.246; Baseline vs APV p=0.000848 RM ANOVA, Tukey Test **(E)** Relative to Baseline (red), NE (gray) shifts action potential firing time closer or ahead of the peak network burst and persists after blocking NMDAR (green). Firing time relative to peak: Baseline= 0.136 ± 0.00817; NE= 0.0616 ± 0.00518; APV= 0.06488 ±0.00397; Baseline vs NE p=<0.0001; NE vs APV p=0.867; Baseline vs APV p=<0.0001. RM ANOVA Tukey Test. Data in A, B and C represent exemplary MEA recordings from a preBötC network. Data in D & E represent n=74 inspiratory units recorded across n=7 networks/slices.

These data motivated us to evaluate changes in the percentage of preBötC network bursts that units participate in during baseline (ACSF) conditions, NE modulation and following blocking NMDAR with APV; these data are represented as percent (%) participation (**Fig. 9A**). On average, the percentage of preBötC network bursts that inspiratory units participated in (% participation) under baseline conditions versus during NE modulation was not significantly different **(Fig. 9A)**.

**Fig. 9.**
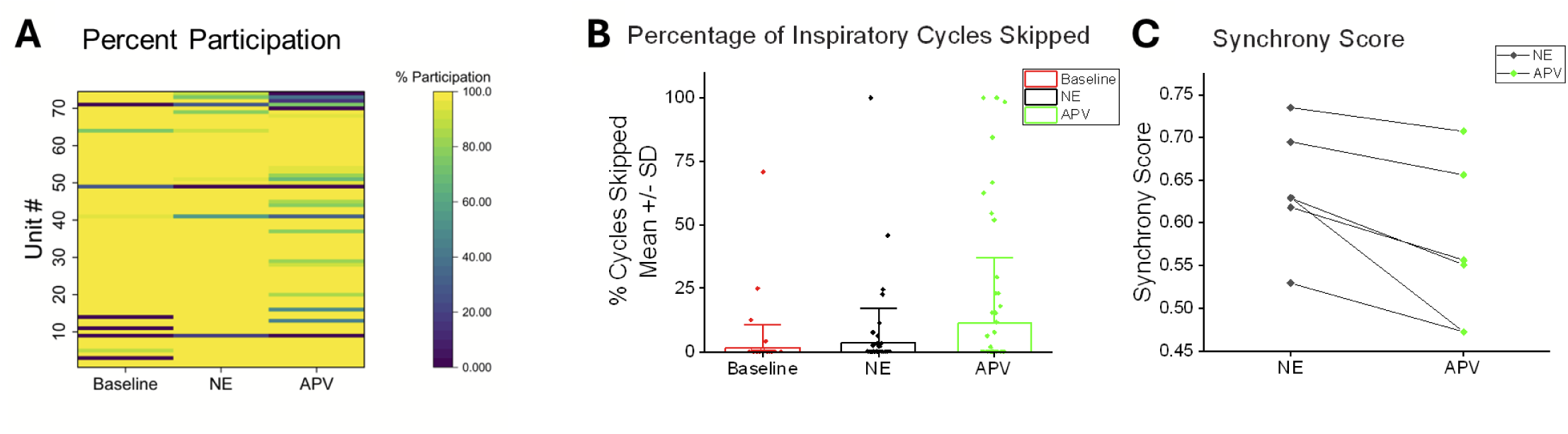
During NE modulation, blockade of NMDAR activity reduces the percentage of preBötC network bursts that inspiratory units participate in, increases the number of preBötC network burst cycles they skip and reduces the overall synchrony score of the preBötC network. **(A)** Heat map of individual participation rates of inspiratory units (n=74 units) during Baseline, NE, and APV conditions showing the heterogeneous effects of NE modulation and APV on activity during the network burst. While the mean participation rate was unchanged in NE (Baseline vs. NE: 91.72 ± 3.07% vs. 94.71% ± 2.09; P=0.583 ANOVA, Tukey Test), it was reduced with APV (APV: 87.57% ± 3.31; Baseline vs. APV p= 0.354; NE vs. APV, p=0.0494, ANOVA, Tukey Test). **(B)** While NE (black) did not increase cycle skipping among inspiratory units participating with the network burst during Baseline conditions (Baseline vs. NE: 1.63 ± 1.10% vs. 3.33% ± 1.65 SE; p=0.738. ANOVA,Tukey test), subsequent blockade of NMDAR by APV (green) increased cycle skipping (APV =11.17% ±3.10; Baseline vs. APV: p=0.000161 Tukey test; NE vs APV p=0.00234 Tukey test). **(C)** During NE modulation, subsequently blocking NMDAR with APV reduces overall network synchrony (mean NE Synchrony Score = 0.639 ±0.0288 SE vs. mean APV score = 0.564±0.03901 SE, p=0.00686, paired t-test).

Across the (n=7) preBötC networks (and n=74 inspiratory units) recorded, NE modulation did not significantly change the likelihood of cycle skipping among units that are active during baseline conditions **(Fig. 9B)**. However, NE recruited five units that were previously silent under baseline conditions into participating in preBötC network bursts (three becoming high-fidelity (HF) units and two low-fidelity (LF) units) and NE increased the firing frequency in the majority (75.7%; n=56/74) of inspiratory units **(Fig. 7E)**. This recruitment and enhanced excitability were in part balanced by reduced excitability of some units, as NE reduced the firing frequency in 24.3% of inspiratory units and silenced others (n=18/74)**(Fig. 7E)**. Further, the response to NE modulation involved changes in unit fidelity whereby four LF units were recruited to become HF and five HF units were de-recruited and became LF units.

To further assess the role of enhanced NMDAR activity in unit fidelity and potential influence of NMDAR activity on individual unit cycle skipping following NE modulation, we applied APV to block NMDAR activity. This NMDAR blockade reduced inspiratory unit fidelity, causing 15 HF inspiratory units to become LF, and silencing three LF inspiratory units. Interestingly, two LF inspiratory units became HF in APV, suggesting complex interactions. After NE modulation, subsequently blocking NMDAR activity reduces the percentage of preBötC network burst cycles inspiratory units participate in **(Fig. 9A)** and increases the probability of cycle skipping **(Fig. 9B)**. During NE modulation, subsequently blocking NMDAR also reduces the synchrony score of the networks (supporting our *in silico* model prediction of the role of NMDAR in synchrony) (**Fig. 9C**).

## Discussion

The primary finding of this study shows that norepinephrine (NE) signaling enhances preBötC network burst amplitude and frequency via recruitment of NMDAR activity. While prior studies in the preBötC have largely identified NMDAR activity as being unnecessary for rhythmogenesis (Connelly et al. 1992; Funk et al. 1997; Lieske and Ramirez 2006; Pace et al. 2007), our data indicates that NMDAR activity is central to shaping network interactions during NE neuromodulation.

Our study dissects how noradrenergic neuromodulation reshapes neuronal activity in the preBötC, revealing that NE-dependent synaptic remodeling is a critical aspect for sculpting and tuning the temporal structure of neuronal interactions during heightened excitability. We identified a potential influence of NE modulation on synaptic inhibition and identified NMDAR-dependent recruitment and shifts in timing of inspiratory neuron activity as central mechanisms through which NE facilitates an increase in preBötC network frequency and strengthening of inspiratory drive. Together, these observations indicate that NE actions on network behavior are not solely by scaling excitability, but rather NE orchestrates coordinated changes in synaptic and intrinsic properties to establish distinct network states capable of generating robust inspiratory drive from the preBötC. While such a coordination can afford stability to network activity, perturbations to either synaptic or intrinsic mechanisms that NE acts upon may promote maladaptive network states and instability in breathing.

In several conditions (Zanella et al. 2014; Browe et al. 2023; Garcia et al. 2016; Kallurkar; et al. 2020) disrupted coordination of action potential activity can lead to subnetwork activity in the preBötC and has downstream consequences, potentially uncoupling the ability of the rhythm generating network to drive downstream motor output, raising the possibility that NMDAR activity is required for maintaining the appropriate E/I balance required for coherent network output and successful transmission of the preBötC rhythm to the motor neuron pool. However, while blockade of NMDAR during NE modulation reduced network burst amplitude, these smaller network events did not appear to be subnetwork bursting, as the preBötC rhythm reliably drove hypoglossal motor output.

While enhanced excitability during NE is primarily attributed to modulation of intrinsic membrane conductances, including evoking conditional pacemaking (Viemari and Ramirez 2006; Venkatakrishnan et al. 2026 (in press)), our findings extend this framework by showing that NE also engages synaptic mechanisms to reshape network dynamics. NMDAR activity enhances the network burst by increasing the degree of temporal alignment of action potentials across the neuronal population during network events. However, after NE modulation, NMDAR blockade reduces both the network synchrony score and cycle-to-cycle participation of inspiratory neurons, underscoring its role in maintaining the fidelity of unit recruitment during the NE-driven state of increased excitability. Importantly, NMDAR recruitment was not a trivial consequence of a shift toward increased excitability as NE modulation was capable of recruiting NMDAR activity even during the disinhibited conditions. Thus, NE-dependent signaling appears to gate NMDAR activity in order to enhance inspiratory rhythmogenesis and facilitate synchronization during neuromodulation.

While NE enhances network excitability and activates NMDAR, subsequent loss of NMDAR function paradoxically accelerates inspiratory burst frequency, while network burst amplitude decreases. The dissociation between rhythm frequency and burst strength highlights the interplay between synaptic excitation, synaptic inhibition and intrinsic properties. Consistent with the concept that enhancing phasic synaptic inhibition during inspiration can drive faster rhythms (Cregg et al. 2017; Baertsch et al. 2018), we found that that disinhibition of the NE-modulated network causes the inspiratory network rhythm to decelerate (Fig. 3), likely reflecting a prolongation of the network refractory period following larger amplitude bursts (Baertsch et al. 2018). These observations implicate a role for synaptic inhibition in NE modulation, facilitating an increase in burst frequency.

Our modeling activities further implicate NMDAR as a central regulator of rhythm frequency when network excitability is elevated. In the CaN network model, where NMDAR are permeable to Ca²⁺ and have slow rise and decay kinetics, increased NMDAR activity prolongs elevations in intracellular Ca²⁺ ([Ca²⁺]ᵢ), that promotes slow ICAN activation and lowers the inactivation threshold of the persistent Na⁺ current (hINaP). Together, these processes slow the network rhythm when NMDAR activity is high. Conversely, reducing NMDAR activity shortens the Ca²⁺ transient, which consequently limits ICAN activation and hINaP inactivation to allow the network to cycle faster, despite the decline in burst amplitude and synchronization. These phenomena were absent in the CaV or CaK models, where the NMDAR is impermeable to Ca²⁺, supporting the importance of NMDAR-mediated Ca^2+^ influx as a rate-limiting Ca²⁺-dependent brake on network frequency during NE-mediated drive. Together, our experiments suggest that frequency modulation by NE is a complex process derived from the integration of synaptic inhibition, synaptic excitation and intrinsic membrane properties.

Prior modeling and experimental work emphasized the importance of synaptically triggered elevations in [Ca²⁺]_i_ (SynCa²⁺) as a driver of ICAN and enhanced preBötC bursting (Phillips et al. 2019). While AMPA receptors are Ca²⁺ impermeable due to GluR2 subunit expression, it has been proposed that kainate receptors, containing the GluK3 subunit are Ca²⁺ permeable and may serve as the principal source of SynCa^2+^ (Phillips et al. 2019). Our combined experimental and in silico data indicate that NMDAR are a critical source of SynCa²⁺ during the NE-modulated state. In the presence of NE, NMDAR activation contributes substantially to both the IDP and the IDC, and subsequent NMDAR blockade returns both measures of inspiratory drive toward their respective baseline levels despite leaving NE-dependent non-NMDAR signaling intact. Our *in silico* modeling efforts also demonstrate that Ca²⁺ permeability of NMDAR is critical for driving bursting that subsequently enhances network burst amplitude and synchrony. Thus, while NMDAR may play a limited role in SynCa^2+^ under standard baseline experimental conditions, our data suggest it likely becomes a dominant source of SynCa²⁺ and determinant of inspiratory network burst amplitude during NE neuromodulation.

The shift of the action potential density of individual units toward the peak of network burst in NE suggests that the neuromodulator drives a network state where NMDAR activity broadens the window of coincidence between presynaptic and postsynaptic activities. This increased coincidence window enhances temporal and spatial summation of synaptic inputs during the network burst. This mechanism may be developmentally and contextually dependent. NMDAR expression and subunit composition changes with age, are modified by synaptic activity (Liu and Wong-Riley; 2010), and may be altered by other neuromodulators (e.g., opioids) and disease states (e.g., intermittent hypoxia and sleep apnea). Thus, the relationship between NE signaling and NMDAR activity may be dynamically tuned across life and during pathology, with implications for how neuromodulatory state governs inspiratory network output and vulnerability to instability. Thus, this study opens the potential for future investigations to take aim at preserving or restoring neuromodulator-dependent synaptic coordination rather than simply scaling synaptic excitability.

In summary, our study demonstrates that NE establishes distinct network states in the preBötC through a coordinated modulation of synaptic and intrinsic neuronal properties, where NMDAR activity plays a central role in shaping both the timing of inspiratory activity across the inspiratory drive. By identifying, NMDAR-dependent Ca^2+^ signaling as a critical determinant of burst amplitude, synchrony and frequency regulation during neuromodulation, we advance a framework where network state during neuromodulation emerges from dynamic interactions between E/I balance, synaptic timing and intrinsic properties. We propose network synchronization is an active process that can be coordinated by neuromodulatory state rather than just being a passive consequence of increased excitability. Thus, this work provides a foundation for understanding how dysregulated neuromodulatory tone may precipitate pathophysiological states with implications for respiratory failure, opioid overdose risk and adaptive limits to homeostatic control.

---

**Supplemental Figure 1.**
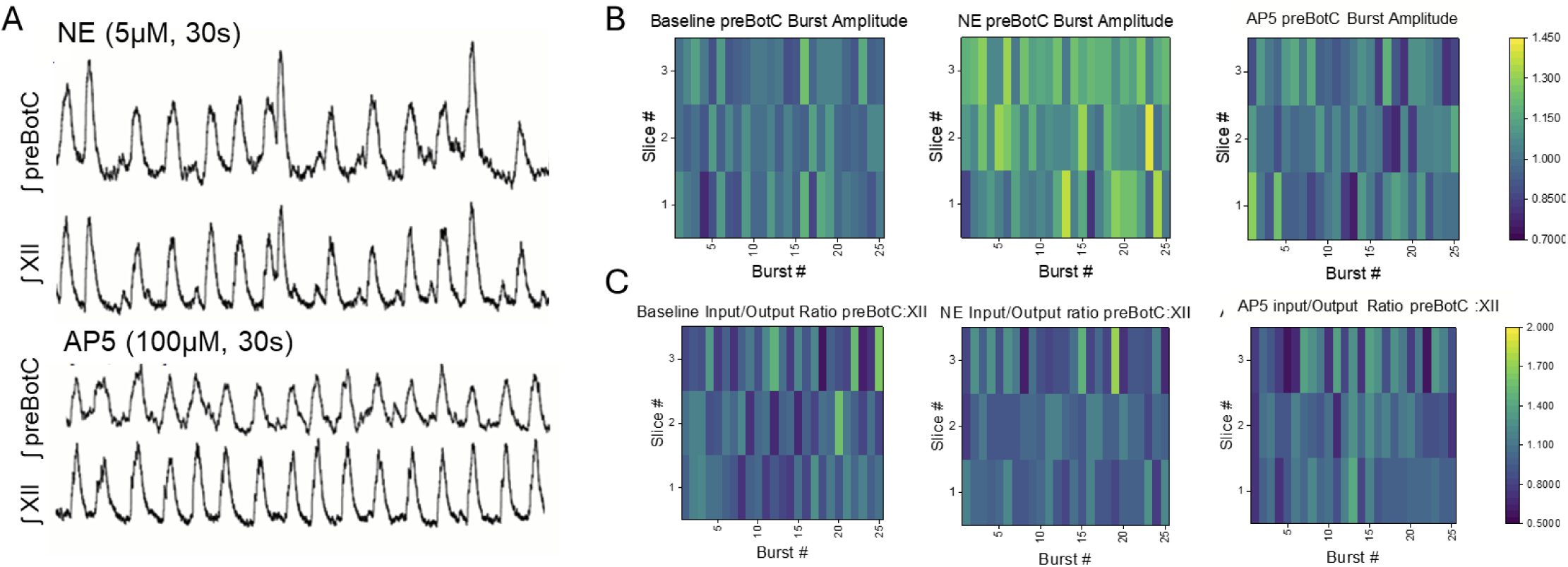
After NE modulation, subsequently blocking NMDAR does not result in loss of the ability of the preBötC to faithfully drive bursting in the hypoglossal (XII) motor pool. (Suppl. **A)** Raw data and **(Suppl. B)** heatmaps illustrating that compared to baseline (ACSF), following bath-application of 5uM NE, integrated preBötC network amplitude (p=0.027 preBötC, One-Sample T-test) and frequency increases (p=0.029, n=3 preBötC; also see Figs 1&2). **(Suppl. A&C)** Following NE-modulation, subsequent blockade of NMDAR with APV does not prevent transmission of preBötC bursting to trigger XII motor output. Heatmap illustrates the input-output relationship between preBötC activity and XII motor output. Heatmaps include n=3 slice preparations (slices # 1-3, y-axis) and 25 consecutive bursts (burst #1-25, x-axis).

**Supplementary Figure 2.**
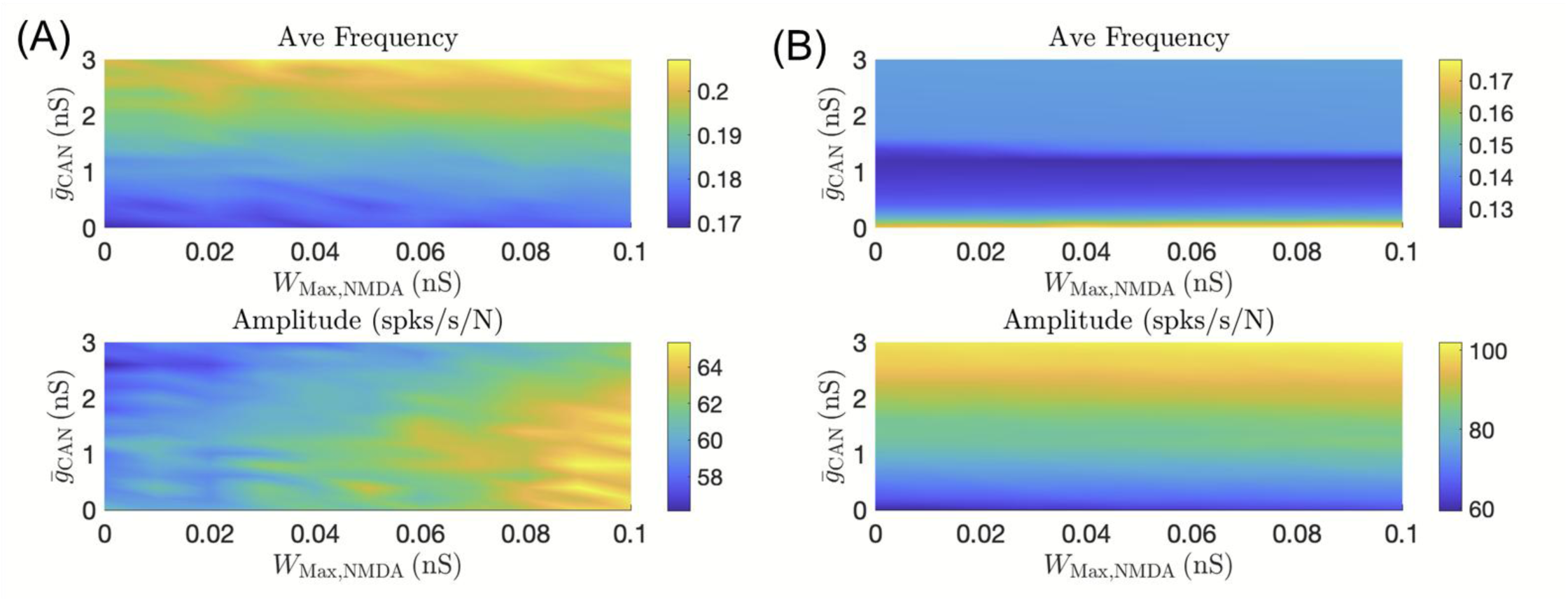
Effects of varying CAN conductance (g ̅_CAN) and NMDAR weight (W_Max,_NMDA) on the dynamics of **(A)** the CaV network where calcium influx is sourced only from voltage-gated calcium channel currents, and **(B)** the CaK network where calcium influx is exclusively from non-NMDA synaptic currents. In both models, for a fixed g ̅_CAN, varryng NMDAR weight has minimal impact on network frequency and amplitude (note the small scales on the color bar for the bottom figure in **(A)**).

