## Supplementary Materials for "Noradrenergic neuromodulation produces a NMDAR-dependent network state of respiratory rhythmogenesis in the preBötzinger Complex"

### 1 Model description

The model describes a network of $N = 100$ synaptically coupled excitatory neurons. For each preBötC neuron, we consider a single-compartment model neuron that incorporates Hodgkin-Huxley-style conductances adapated from previously described models (Jasinski et al., 2013; Wang and Rubin, 2017; Phillips et al., 2019; Phillips and Rubin, 2022). Each neuron's dynamics are governed by the following differential equations:

$$
\begin{aligned}
C\frac{dV}{dt} &= -I_{\mathrm{Na}} - I_{\mathrm{NaP}} - I_{\mathrm{K}} - I_{\mathrm{Ca}} - I_{\mathrm{CAN}} - I_{\mathrm{L}} - I_{\mathrm{NMDA}} - I_{\mathrm{nonNMDA}} - I_{\mathrm{Tonic}} \\
\frac{dy}{dt} &= (y_{\infty}(V) - y)/\tau_y(V), \quad y = \{m_{\mathrm{Na}}, h_{\mathrm{Na}}, m_{\mathrm{Nap}}, h_{\mathrm{Nap}}, m_{\mathrm{Ca}}, h_{\mathrm{Ca}}, m_{\mathrm{K}}\} \\
\frac{d\mathrm{Ca_{tot}}}{dt} &= -\alpha_{\mathrm{Ca}}(P_{\mathrm{ICa}} I_{\mathrm{Ca}} + P_{\mathrm{nonNMDAca}} \cdot I_{\mathrm{nonNMDA}} + P_{\mathrm{NMDAca}} \cdot I_{\mathrm{NMDA}}) \\
&\quad -(\mathrm{Ca_i} - \mathrm{Ca_{min}})/\tau_{\mathrm{Ca}} \\
\frac{d\mathrm{Ca_i}}{dt} &= -\alpha_{\mathrm{Ca}}(P_{\mathrm{ICa}} I_{\mathrm{Ca}} + P_{\mathrm{nonNMDAca}} \cdot I_{\mathrm{nonNMDA}} + P_{\mathrm{NMDAca}} \cdot I_{\mathrm{NMDA}}) \\
&\quad +\alpha_{\mathrm{ER}}(J_{\mathrm{ER_{IN}}} - J_{\mathrm{ER_{OUT}}}) - (\mathrm{Ca_i} - \mathrm{Ca_{min}})/\tau_{\mathrm{Ca}} \\
\frac{dl}{dt} &= AK_{\mathrm{d}}(1-l) - A\mathrm{Ca_i} l
\end{aligned}
\tag{1.1}
$$

The neuronal membrane potential ($V$) is governed by a set of membrane ionic currents. $C = 36\,\mathrm{pF}$ is neuronal membrane capacitance and $t$ is time. Each $I_i$ represents a current with $i$ denoting the current's type. The ionic currents in the model include fast $Na^+$ current ($I_{\mathrm{Na}}$), persistent $Na^+$ current ($I_{\mathrm{NaP}}$), delayed rectifier $K^+$ current ($I_{\mathrm{K}}$), high-voltage-activated $Ca^{2+}$ current ($I_{\mathrm{Ca}}$), $Ca^{2+}$-activated nonspecific cation current ($I_{\mathrm{CAN}}$), $K^+$-dominated leak current ($I_{\mathrm{L}}$), excitatory synaptic currents ($I_{\mathrm{nonNMDA}}$ and $I_{\mathrm{NMDA}}$). $I_{\mathrm{Tonic}}$ denotes a fixed or tonic excitatory synaptic input (e.g., from respiratory control areas outside of the preBotC). The currents are defined in Table 1.

In Table 1, $g_i$ is the maximum conductance. Activation ($m_i$) and inactivation ($h_i$) variables for most ionic channels $I_i$ are governed by the second $y$ equation in (1.1), where steady-state activation ($m_{\infty}$) and inactivation ($h_{\infty}$) functions and time constants are described as in Table 2. Notice from the bottom row in Table 2 that unlike the other currents, $I_{\mathrm{CAN}}$ has

Table 1: Ionic currents and channel reversal potentials for system (1.1)

| Currents (pA) | Reversal potentials (mV) |
|---|---|
| $I_{\mathrm{Na}} = g_{\mathrm{Na}} m_{\mathrm{Na}}^3 h_{\mathrm{Na}} (V - E_{\mathrm{Na}})$ <br> $I_{\mathrm{NaP}} = g_{\mathrm{NaP}} m_{\mathrm{NaP}} h_{\mathrm{NaP}} (V - E_{\mathrm{Na}})$ | $E_{\mathrm{Na}} = 26.54 \ln(Na_o/Na_i)$ |
| $I_{\mathrm{K}} = g_{\mathrm{K}} m_{\mathrm{K}}^4 (V - E_{\mathrm{K}})$ | $E_{\mathrm{K}} = 26.54 \ln(K_o/K_i)$ |
| $I_{\mathrm{Ca}} = g_{\mathrm{Ca}} m_{\mathrm{Ca}} h_{\mathrm{Ca}} (V - E_{\mathrm{Ca}})$ | $E_{\mathrm{Ca}} = 13.27 \ln(\mathrm{Ca_o}/\mathrm{Ca_i})$ |
| $I_{\mathrm{CAN}} = g_{\mathrm{CAN}} m_{\mathrm{CAN}} (V - E_{\mathrm{CAN}})$ | $E_{\mathrm{CAN}} = 0$ |
| $I_{\mathrm{L}} = g_{\mathrm{L}} (V - E_{\mathrm{L}})$ | $E_{\mathrm{L}} = -26.54 \ln((Na_i + 42K_i)/(Na_o + 42K_o))$ |
| $I_{\mathrm{nonNMDA}} = g_{\mathrm{nonNMDA}} (V - E_{\mathrm{SynE}})$ <br> $I_{\mathrm{NMDA}} = g_{\mathrm{NMDA}} B(V) (V - E_{\mathrm{SynE}})$ <br> $I_{\mathrm{Tonic}} = g_{\mathrm{Tonic}} (V - E_{\mathrm{SynE}})$ | $E_{\mathrm{SynE}} = 0$ |

instantaneous activation, which depends on $\mathrm{Ca_i}$ and is voltage-independent. The parameters for these currents are specified in Table 3.

Table 2: Functions associated with activation and inactivation variables for system (1.1). We use the variable $y$ when an expression corresponds to a set of variables.

| gating variables | steady state activation and inactivation | time constants |
|---|---|---|
| $m_{\mathrm{Na}}$ | $y_\infty(V) = 1/(1 + \exp(-(V - V_{y1/2})/k_y))$ | $\tau_y(V) = \tau_{y\max}/\cosh(-(V - V_{\tau y1/2})/k_{\tau y})$ |
| $h_{\mathrm{Na}}$ | | |
| $m_{\mathrm{NaP}}$ | | |
| $h_{\mathrm{Na}}$ | | |
| $m_{\mathrm{Ca}}$ | | $\tau_{m_{\mathrm{Ca}}} = 0.5\,\mathrm{ms}$ |
| $h_{\mathrm{Ca}}$ | | $\tau_{h_{\mathrm{Ca}}} = 18\,\mathrm{ms}$ |
| $m_{\mathrm{K}}$ | $m_{\mathrm{K}\infty} = \alpha_\infty/(\alpha_\infty + \beta_\infty)$ | $\tau_{m_{\mathrm{K}}} = 1/(\alpha_\infty + \beta_\infty)$ |
| | $\alpha_\infty = A_\alpha \cdot (V + B_\alpha)/(1 - \exp(-(V + B_\alpha)/k_\alpha))$, $\beta_\infty = A_\beta \cdot \exp(-(V + B_\beta)/k_\beta)$ | |
| $m_{\mathrm{CAN}}$ | $m_{\mathrm{CAN}} = 1/(1 + (K_{\mathrm{CAN}}/\mathrm{Ca_i})^n)$ | |
| Magnesium block | $B(V) = 1/(1 + e^{-0.062V}[\mathrm{Mg}^{2+}]/3.57)$ | |

The dynamics of the total intracellular $\mathrm{Ca}^{2+}$ concentration within the cell ($\mathrm{Ca_{tot}}$) and intracellular concentration of free $\mathrm{Ca}^{2+}$ ($\mathrm{Ca_i}$) are described by the third and fourth equations in (1.1), respectively. Both variables are intimately linked with $l$, the fraction of $\mathrm{IP}_3$ receptors that are not inactivated by calcium, which is governed by the last equation of (1.1). In the calcium equations, $J_{\mathrm{ERin}}$ represents the flux of $\mathrm{Ca}^{2+}$ per unit volume from the endoplasmic reticulum (ER) into the cytoplasm, which depends on $l$, and $J_{\mathrm{ERout}}$ represents the flux of $\mathrm{Ca}^{2+}$ per unit volume from the cytoplasm into the ER. These two fluxes are modeled by (1.2):

Table 3: Parameter values for system (1.1)

| Current | Parameters |
|---|---|
| Fast $Na^+$ ($I_{\mathrm{Na}}$) | $g_{\mathrm{Na}} = 150\,\mathrm{nS}$, $Na_i = 15$ mM, $Na_o = 120$ mM |
| | $V_{m1/2} = -43.8\,\mathrm{mV}, k_m = 6\,\mathrm{mV}, \tau_{mmax} = 0.25\,\mathrm{ms},$ $V_{\tau m1/2} = -43.8\,\mathrm{mV}, k_{\tau m} = 14\,\mathrm{mV}$ $V_{h1/2} = -67.5\,\mathrm{mV}, k_h = -11.8\,\mathrm{mV}$ $\tau_{hmax} = 8.46\,\mathrm{ms}$, $V_{\tau h1/2} = -67.5\,\mathrm{mV}, k_{\tau h} = 12.8\,\mathrm{mV}$ |
| Persistent $Na^+$ ($I_{\mathrm{NaP}}$) | $g_{\mathrm{NaP}} = N(\mu_{\mathrm{NaP}}, \sigma_{\mathrm{NaP}})\,\mathrm{nS}$, $\mu_{\mathrm{NaP}} = 3.33\,\mathrm{nS}$ $\sigma_{\mathrm{NaP}} = 0.75\,\mathrm{nS}$ |
| | $V_{m1/2} = -47.1\,\mathrm{mV}, k_m = 3.1\,\mathrm{mV}, \tau_{mmax} = 1\,\mathrm{ms},$ $V_{\tau m1/2} = -47.1\,\mathrm{mV}, k_{\tau m} = 6.2\,\mathrm{mV}$ $V_{h1/2} = -60\,\mathrm{mV}, k_h = -9\,\mathrm{mV}, \tau_{hmax} = 5000\,\mathrm{ms},$ $V_{\tau h1/2} = -60\,\mathrm{mV}, k_{\tau h} = 9\,\mathrm{mV}$ |
| $K^+$ delayed rectifier ($I_K$) | $g_K = 220\,\mathrm{nS}$, $K_o = 6.5$ mM, $K_i = 125$ mM |
| | $A_\alpha = 0.01, B_\alpha = 44\,\mathrm{mV}, k_\alpha = 5\,\mathrm{mV}, A_\beta = 0.17,$ $B_\beta = 49\,\mathrm{mV}, k_\beta = 40\,\mathrm{mV}$ |
| $Ca^{2+}$ ($I_{\mathrm{Ca}}$) | $g_{\mathrm{Ca}} = 0.000065\,\mathrm{nS}$, $\mathrm{Ca_o} = 4$ mM |
| | $V_{m1/2} = -27.5\,\mathrm{mV}, k_m = 5.7\,\mathrm{mV}$ $V_{h1/2} = -52.4\,\mathrm{mV}, k_h = -5.2\,\mathrm{mV}$ |
| $Ca^{2+}$-activated nonspecific ($I_{\mathrm{CAN}}$) | $g_{\mathrm{CAN}} = \mathrm{VAR}, K_{\mathrm{CAN}} = 0.00074\,\mathrm{mM}, n = 0.97$ |
| Leakage ($I_L$) | $g_L = N(\mu_L, \sigma_L)$, $\mu_L = exp((K_o - 3.425)/4.05)\,\mathrm{nS}$ $\sigma_L = 0.05 \cdot \mu_L\,\mathrm{nS}$ |
| non-NMDA synaptic ($I_{\mathrm{nonNMDA}}$) | $g_{\mathrm{nonNMDA}} = \mathrm{VAR}$, see (1.3), $\tau_{\mathrm{nonNMDA}} = 5\,\mathrm{ms}$ |
| NMDA synaptic ($I_{\mathrm{NMDA}}$) | $g_{\mathrm{NMDA}} = \mathrm{VAR}$, see (1.3), $\tau_{\mathrm{NMDA}} = 20\,\mathrm{ms}$, $[\mathrm{Mg}^{2+}] = 2\,\mathrm{mM}$ |
| Tonic synaptic ($I_{\mathrm{Tonic}}$) | $g_{\mathrm{Tonic}} = 0.3\,\mathrm{nS}$ |

$$J_{\mathrm{ER_{IN}}} = \left( L_{\mathrm{IP_3}} + P_{\mathrm{IP_3}} \left[ \frac{[\mathrm{IP_3}]\mathrm{Ca_i} l}{([\mathrm{IP_3}] + K_I)(\mathrm{Ca_i} + K_a)} \right]^3 \right) (\mathrm{Ca_{ER}} - \mathrm{Ca_i}) \tag{1.2a}$$

$$J_{\mathrm{ER_{OUT}}} = V_{\mathrm{SERCA}} \frac{\mathrm{Ca_i^2}}{K_{\mathrm{SERCA}}^2 + \mathrm{Ca_i^2}} \tag{1.2b}$$

$$\mathrm{Ca_{ER}} = \frac{\mathrm{Ca_{tot}} - \mathrm{Ca_i}}{\sigma}. \tag{1.2c}$$

The values of parameters associated with the equations for $\mathrm{Ca_{tot}}$, $\mathrm{Ca_i}$ and $l$ are given by: $\alpha_{\mathrm{Ca}} = 2.5 \times 10^{-5}\,\mathrm{mM/fC}$, $\tau_{\mathrm{Ca}} = 500$ ms, $\mathrm{Ca_{min}} = 5.0 \times 10^{-6}\,\mathrm{mM}$, $\alpha_{\mathrm{ER}} = 2.5 \times 10^{-5}$, $L_{\mathrm{IP3}} = 0.1/\mathrm{ms}$, $P_{\mathrm{IP_3}} = 77500/\mathrm{ms}$, $[\mathrm{IP_3}] = 1.5 \times 10^{-3}\,\mathrm{mM}$, $K_I = 10^{-3}\,\mathrm{mM}$, $K_a = 10^{-4}\,\mathrm{mM}$, $V_{\mathrm{SERCA}} = 0.45\,\mathrm{mM/ms}$, $K_{\mathrm{SERCA}} = 7.5 \times 10^{-5}\,\mathrm{mM}$, $\sigma = 0.185$, $A = 0.1\,\mathrm{mM/ms}$ and $K_{\mathrm{d}} = 0.2 \times 10^{-3}\,\mathrm{mM}$. Additional description of model components has been given previously Toporikova and Butera (2011); Park and Rubin (2013); Jasinski et al. (2013); Wang and Rubin (2017); Phillips and Rubin (2022).

In addition, there are various sources of $Ca^{2+}$ influx from the extracellular space: voltage-gated $Ca^{2+}$ channels ($I_{\mathrm{Ca}}$) and synaptic inputs where a percentage of the synaptic current is assumed to consist of $Ca^{2+}$ ions. The specific mechanisms behind the synaptic calcium sources are unclear and likely depend on specific types of glutamate receptors. There are three subtypes of ionotropic glutamate receptors, N-methyl-D-aspartate (NMDA), Kainate (KAR), and a-amino-3-hydroxy-5-methyl-4-isoxazolepropionic acid (AMPA), all of which are expressed in the pre-BötC and have varying degrees of calcium permeability. Among them, AMPA is not permeable to $Ca^{2+}$, whereas KAR and NMDA are possible candidates for synaptically mediated calcium entry (Phillips et al., 2019; Morgado-Valle and Feldman, 2007; Ermentrout and Terman, 2010). As a result, we include three possible sources for calcium influx in the $\mathrm{Ca_i}$ and $\mathrm{Ca_{tot}}$ equations (see (1.1)): $P_{\mathrm{ICa}} I_{\mathrm{Ca}}$, $P_{\mathrm{nonNMDAca}} \cdot I_{\mathrm{nonNMDA}}$ and $P_{\mathrm{NMDAca}} \cdot I_{\mathrm{NMDA}}$. The parameters $P_{\mathrm{ICa}}$, $P_{\mathrm{nonNMDAca}}$ and $P_{\mathrm{NMDAca}}$ control the relative calcium permeability from each source. Following (Phillips et al., 2019), we denote the model with $P_{\mathrm{nonNMDAca}} = P_{\mathrm{NMDAca}} = 0$ as the CaV model, the model with $P_{\mathrm{ICa}} = P_{\mathrm{NMDAca}} = 0$ as the CaK model, and the model with $P_{\mathrm{ICa}} = P_{\mathrm{nonNMDAca}} = 0$ as the CaN model.

In (1.1), the synaptic conductance of fast AMPA/kainate synapse (denoted as $g^i_{\mathrm{nonNMDA}}$) and NMDA receptors ($g^i_{\mathrm{NMDA}}$) for the $i$-th neuron in the population are described by the following equations:

$$\begin{aligned} g^i_{\mathrm{nonNMDA}} &= \textstyle\sum_{j \neq i,n} W_{\mathrm{nonNMDA},ji} D_j C_{j,i} H(t - t_{j,n}) e^{-(t-t_{j,n})/\tau_{\mathrm{nonNMDA}}} \\ g^i_{\mathrm{NMDA}} &= \textstyle\sum_{j \neq i,n} W_{\mathrm{NMDA},ji} D_j C_{ji} H(t - t_{j,n}) e^{-(t-t_{j,n})/\tau_{\mathrm{NMDA}}} \end{aligned} \tag{1.3}$$

where $W_{\mathrm{nonNMDA},ji}$ and $W_{\mathrm{NMDA},ji}$ represent the weights of the non-NMDA and NMDA synaptic connection from neuron $j$ to neuron $i$. $D_j$ is a scaling factor for short-term synaptic depression in the presynaptic neuron $j$. It evolves according to the equation $\frac{dD_j}{dt} = \frac{D_0 - D_j}{\tau_D} - \alpha_D D_j \delta(t - t_j)$, where $D_0 = 1$, $\tau_D = 1000\,\mathrm{ms}$, $\alpha_D = 0.2$ and $\delta(.)$ is the Kronecker delta function that equals 1 at the time of each spike in neuron $j$ and 0 otherwise. Additional description of the synaptic depression and related parameters have been given in (Phillips and Rubin, 2022). $C_{j,i}$ is an element of the connectivity matrix ($C_{j,i} = 1$ if neuron $j$ makes a synapse with neuron $i$ and $C_{j,i} = 0$ otherwise), $H(.)$ is the Heaviside step function, and $t$ denotes time. $\tau_{\mathrm{nonNMDA}}$ and $\tau_{\mathrm{NMDA}}$ are exponential synaptic decay constants for non-NMDA and NMDA receptors (see Table 3). $t_{j,n}$ is the time at which the $n$-th action potential generated by neuron $j$ reaches neuron $i$.

The current for an NMDA synapse is modeled differently from that of AMPA/kainate (see Table 1) because, under normal physiological conditions, the NMDA receptor is partially blocked by magnesium ions (Ermentrout and Terman, 2010). Specificically, $B(V)$, as defined in the last row of Table 2 within the $I_{\mathrm{NMDA}}$ function in Table 1, represents this magnesium block. As discussed above, these currents contribute to synaptically triggered $Ca^{2+}$ transients (see (1.1)), where $P_{\mathrm{nonNMDAca}}$ and $P_{\mathrm{NMDAca}}$) denote the percentages of $Ca^{2+}$ ions that can pass through AMPA/kainate ($I_{\mathrm{nonNMDA}}$) and NMDA synaptic currents ($I_{\mathrm{NMDA}}$), respectively. In our simulations, the calcium permeability parameters were varied over the range $P_{\mathrm{nonNMDAca}} \in [0, 0.0015]$ and $P_{\mathrm{NMDAca}} \in [0, 0.005]$. Excitatory synaptic weights were drawn from a uniform distribution, $W_{j,i} = U(0, W_{\mathrm{Max}})$, with $W_{\mathrm{Max}} = 0.15\,\mathrm{nS}$ for non-NMDA receptors and $W_{\text{Max}} = 0.05\,\text{nS}$ for NMDA receptors. The higher range for $P_{\text{NMDAca}}$ reflects the relatively higher calcium permeability of NMDARs, whereas calcium influx through non-NMDA receptors primarily arises from KARs Phillips et al. (2019); Paarmann et al. (2000); Perrais et al. (2009). In contrast, the smaller maximum NMDAR synaptic weight reflects experimental findings that, under normal conditions, NMDAR-mediated coupling is significantly weaker than that of AMPARs in the preBotC (Morgado-Valle and Feldman, 2007). In addition, $g_{\text{CAN}}$ was uniformly distributed over the range $[0, \bar{g}_{\text{CAN}}]$ nS. The elements of the network connectivity matrix, $C_{j,i}$, are randomly assigned values of 0 or 1 such that the probability of any connection between neuron $j$ and neuron $i$ being 1 is equal to the network connection probability $P_{\text{Syn}} = 0.13$. We examine the effects of NMDA synaptic strength and $I_{\text{CAN}}$ on the dynamcis of our model by varying $W_{\text{Max,NMDA}}$ and $\bar{g}_{\text{CAN}}$ (see, e.g., supplentary Figure 2).
